# Nicotinamide provides neuroprotection in glaucoma by protecting against mitochondrial and metabolic dysfunction

**DOI:** 10.1101/2020.10.21.348250

**Authors:** James R Tribble, Amin Otmani, Shanshan Sun, Sevannah A Ellis, Gloria Cimaglia, Rupali Vohra, Melissa Jöe, Emma Lardner, Abinaya P Venkataraman, Alberto Domínguez-Vicent, Eirini Kokkali, Seungsoo Rho, Gauti Jóhannesson, Robert W Burgess, Peter G Fuerst, Rune Brautaset, Miriam Kolko, James E Morgan, Jonathan G Crowston, Marcela Votruba, Pete A Williams

**Author notes:** To whom correspondence should be addressed: Pete A Williams; Department of Clinical Neuroscience, Division of Eye and Vision, St. Erik Eye Hospital, Karolinska Institutet, Stockholm, Sweden.

## Abstract

Nicotinamide adenine dinucleotide (NAD) is a REDOX cofactor and metabolite essential for neuronal survival. Glaucoma is a common neurodegenerative disease in which neuronal levels of NAD decline. Repleting NAD via dietary supplementation of nicotinamide (a precursor to NAD) is effective in preventing retinal ganglion cell neurodegeneration in mouse models. Supporting this, short-term oral nicotinamide treatment in human glaucoma patients provides a recovery of retinal ganglion cell function implying a protection of visual function. Despite this, the mechanism of neuroprotection and full effects of nicotinamide on retinal ganglion cells is unclear. Glaucoma is a complex neurodegenerative disease in which a mix of healthy, stressed, and degenerating retinal ganglion cells co-exist, and in which retinal ganglion cells display compartmentalized degeneration across their visual trajectory. Therefore, we assess the effects of nicotinamide on retinal ganglion cells in normal physiological conditions and across a range of glaucoma relevant insults. We confirm neuroprotection afforded by nicotinamide in rodent models which represent isolated ocular hypertensive, axon degenerative, and mitochondrial degenerative insults. We define a small molecular weight metabolome for the retina, optic nerve, and superior colliculus which demonstrates that ocular hypertension induces widespread metabolic disruption that can be prevented by nicotinamide. Nicotinamide provides these neuroprotective effects by increasing oxidative phosphorylation, buffering and preventing metabolic stress, and increasing mitochondrial size and motility whilst simultaneously dampening action potential firing frequency. These data support continued determination of the utility of long-term NAM treatment as a neuroprotective therapy for human glaucoma.

**One Sentence Summary:** The NAD precursor nicotinamide has a potent neuroprotective effect in the retina and optic nerve, targeting neuronal function, metabolism, and mitochondrial function.

## Introduction

Glaucoma is one of the most common neurodegenerative diseases, and the leading cause of irreversible blindness, affecting ~80 million people worldwide (1). Age, genetics, and elevated intraocular pressure (IOP) are major risk factors, but to date, the only clinically available therapies target the reduction of IOP. For the many patients which are refractory to IOP lowering treatments, surgical interventions are commonly used in order to limit the progressive neurodegeneration and visual impairment (2). No therapy currently targets the neuronal population that degenerates in glaucoma. Ultimately, 42% of treated glaucoma patients progress to blindness in at least one eye (3). With an increasingly aged population, glaucoma will continue to be a significant health and economic burden and thus neuroprotective strategies targeting the retina and optic nerve are of great therapeutic need.

Retinal ganglion cells, the output neuron of the retina whose axons converge to form the optic nerve, display a progressive, compartmentalized neurodegeneration leading to the characteristic visual dysfunction seen in glaucoma. We have previously demonstrated mitochondrial abnormalities occurring prior to neurodegeneration in glaucoma (in glaucoma patients and animal models (4, 5)). The latter study, utilizing the DBA/2J mouse model of glaucoma, has identified that nicotinamide adenine dinucleotide (NAD; an essential REDOX cofactor and metabolite) declines in the retina in an age-dependent manner and renders retinal ganglion cells susceptible to IOP-related stress, driving glaucomatous neurodegeneration. The prevention of NAD decline by dietary supplementation with nicotinamide (NAM; the amide form of vitamin B3, an NAD precursor via the NAD-salvage pathway in neurons) or by intravitreal viral gene-therapy overexpressing *Nmnat1* (a terminal enzyme for NAD production) was robustly protective against retinal ganglion cell neurodegeneration and prevented a number of early gene expression changes in DBA/2J retinal ganglion cells (5). Glaucoma patients have been demonstrated to have systemically low levels of NAM in sera (6) which supports a hypothesis in which pathogenically low NAD leads to glaucoma susceptibility. These findings have driven interest in nicotinamide as a treatment for glaucoma, including the planning and initiation of a number of clinical trials, *e.g*. NCT03797469 which is ongoing and Hui *et al*. (2020) which is now completed (7). In this trial, as part of a multi-national collaborative team, we have demonstrated significant visual recovery in glaucoma patients over a 6 month treatment period (cross-over design; step up 1.5 g/d to 3.0 g/d nicotinamide (7)). Nicotinamide’s excellent long clinical history, combined with good safety profile, tolerance at high doses, and affordability has facilitated rapid translation into initial small scale clinical trials (8, 9). However, translation to larger scale trials, the evaluation of earlier intervention protocols, and/or longer-term clinical outcomes requires a deeper understanding of the effects of nicotinamide on retinal ganglion cells under both neurodegenerative conditions and normal physiological states. This is of critical importance since disease progression is often heterogeneous (*i.e*. a mix of healthy, degenerating, and dead cells) and if prophylactic treatment is to be considered (which is ideal since visual deficits present once the degenerative processes have already been initiated (10)).

We bridge the gap between the bench and bedside by demonstrating further evidence of neuroprotection through nicotinamide supplementation across a number of glaucoma-related retinal ganglion cell targeted insults, using a number of retinal ganglion cell specific tools. We identify early metabolic changes in the retina in a rat model of ocular hypertensive glaucoma (elevated IOP), which are prevented by nicotinamide supplementation. Importantly, chronic dietary nicotinamide supplementation causes minimal metabolic variation in non-diseased visual system tissues supporting potential prophylactic treatments. Nicotinamide administration also buffers retinal ganglion cell metabolism, increases oxidative phosphorylation, and positively alters mitochondrial morphology and dynamics in retinal ganglion cells at doses which are neuroprotective. These data help support nicotinamide’s use in the clinic, and support the initiation of clinical trials to determine the effect of NAM on longer-term disease progression.

## Results

### Nicotinamide confers retinal ganglion cell neuroprotection in a range of glaucoma-relevant contexts

Nicotinamide is protective in a genetic mouse model (DBA/2J) that recapitulates features of a human pigmentary glaucoma and demonstrates an age-related decline in retinal NAD (5). To further determine how nicotinamide (NAM) protects retinal ganglion cells (RGCs), we induced RGC degeneration in two separate glaucoma relevant insults in the absence of an age related effect: elevated IOP and optic nerve degeneration induced by axotomy. We first confirmed the neuroprotective effects of NAM in an inducible, acute rat model of glaucoma where the onset of high IOP is controlled and abrupt. Restriction of the anterior chamber drainage structures via magnetic microbead injection limits aqueous humor fluid outflow, producing significant ocular hypertension (OHT; elevated intraocular pressure) which is sustained for two weeks (**Figure 1A**). NAD was significantly reduced in the retina and optic nerve of OHT animals at 14 days postinduction (**Figure 1B**) (*i.e*. IOP-dependent, not age-dependent NAD decline). Clinical imaging using optical coherence tomography (OCT) demonstrated loss of the neuroretinal rim (representing intra-retinal retinal ganglion cell axons; **Figure 1C**). Rats pre-treated with NAM (200 mg/kg/d dietary, representative of 2 g/d for a 60 kg human (11); with treatment continuing for the duration of OHT) did not demonstrate loss of the neuroretinal rim despite maintaining a similar IOP (**Figure 1C**). OHT eyes demonstrated significant RGC loss as assessed by RBPMS+ counting (an RGC specific marker in the retina, **Figure 1D**). The average nuclear diameter (DAPI+) was also significantly decreased, suggesting that surviving cells were under considerable neurodegenerative stress (**Figure 1E**) (12). NAM treated OHT rats demonstrated significantly reduced RGC loss and nuclear shrinkage in a dose dependent manner (200, 400, or 800 mg/kg/d dietary, representative of 2, 4, or 8 g/d for a 60 kg human (11); **Figure 1E**). These data support a neuroprotective role for NAM in the context of high IOP without an associated age-related NAD decline.

**Figure 1.**
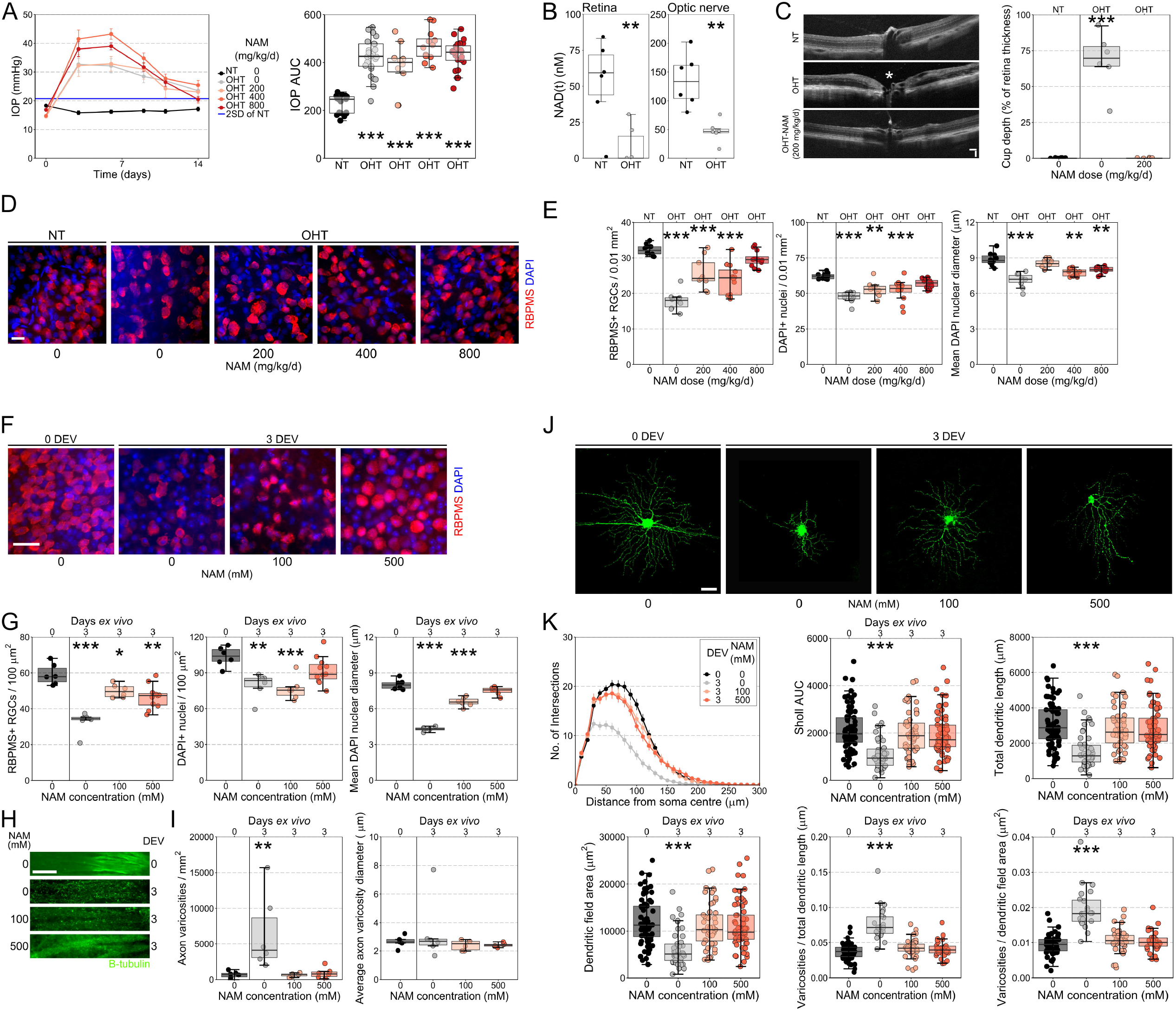
Nicotinamide provides retinal ganglion cell neuroprotection. (**A**) Ocular hypertension (OHT; *n* = 26 eyes) induced by the magnetic bead injection lead to a significant increase in IOP over control (NT; *n* = 20 eyes), which persisted for 14 days (greater than 2 standard deviations of NT, blue line). Nicotinamide (NAM) treatment in OHT animals (200, 400, 800 mg/kg/d; *n* = 10, 12, and 24 eyes respectively) did not lower IOP (as assessed by area under the IOP curve). (**B**) Following 14 days of sustained OHT, NAD was significantly reduced in both the retina (*n* = 6) and optic nerve (*n* = 6) as measured by luminometry-based assays. (**C**) *In vivo* OCT imaging demonstrated significant loss of neuroretinal rim following 14 days OHT (*n* = 7 eyes) compared to NT eyes (*n* = 9), which was absent in NAM treated rats (*n* = 8 eyes; measured as the cup depth relative to retinal thickness; loss denoted by white * on example image). (**D**) RBPMS (RGC specific) and DAPI labelling of the retina demonstrated a dose dependent, significant neuroprotection against RGC loss and nuclear shrinkage at day 14 (**E**; *n* = 10 NT retinas, 10 OHT, 9 OHT-NAM (200 mg/kg/d), 12 OHT-NAM (400 mg/kg/d), and 12 OHT-NAM (800 mg/kg/d)). (**F**) Retinal axotomy explant results in significant RGC death and nuclear shrinkage, at 3 days *ex vivo* (DEV) with robust protection provided by NAM treated media (**G**; *n* = six 0 DEV retinas, six 3 DEV untreated, six 3 DEV-NAM (100 mM), eleven 3 DEV-NAM (500 mM)). (**H**) β-tubulin labelling reveled a significant increase in axonal varicosities within the retinal nerve fiber layer at 3 DEV, which were robustly protected against by NAM (**I**; *n* = six 0 DEV retinas, six 3 DEV untreated, six 3 DEV-NAM (100 mM), six 3 DEV-NAM (500 mM)). (**J and K**) At 3 DEV significant changes to the complexity of the dendritic arbor were apparent (**J**; example RGCs labelled by DiOlistics). At 3 DEV there was a loss of branch density (reduction in Sholl AUC), a reduction in dendritic length and field area. NAM treated media provided significant protection against these neurodegenerative features (*n* = 75 RGCs from ten 0 DEV retinas, 46 RGCs from twelve 3 DEV untreated, 53 RGCs from twelve 3 DEV-NAM (100 mM), 62 RGCs from twelve 3 DEV-NAM (500 mM)). At 3 DEV dendrites display an increased density of varicosities, which was protected against by NAM (*n* = 55 RGCs from 0 DEV retinas, 23 RGCs from 3 DEV untreated, 38 RGCs from 3 DEV-NAM (100 mM), 32 RGCs from 3 DEV-NAM (500 mM)). Scale bar = 100 μm in C, 20 μm in D and F, 50 μm in H and J. * *P* < 0.05, ** *P* < 0.01, *** *P* < 0.001.

We next tested NAM protection in an acute, axon-specific injury using a retinal explant model (13, 14). Enucleation severs RGC axons in the optic nerve, and explants maintained in culture *ex vivo*, demonstrated a progressive RGC loss and nuclear shrinkage observed within 12 hours, with highly significant, reproducible RGC loss at 3 days *ex vivo* (**Figure 1F and G, Supplementary Figure 1A and B**). This RGC loss and nucleus shrinkage is robustly prevented by NAM-supplement media at a range of doses. Using this explant model we assessed whether protection extended to RGC axons and dendrites. RGC axons within the retinal nerve fiber layer (RNFL) demonstrated a significant increase in the number of varicosities, a metric of neurite health and degeneration, by 3 days *ex vivo*, but with no change in average varicosity size (**Figure 1H and I, Supplementary Figure 1C and D**). These changes were not observed in NAM treated retina, even by 3 days *ex vivo*, demonstrating significant protection against severe acute axon degeneration (**Figure 1H and I**). At 3 days *ex vivo* RGCs had significant dendritic atrophy compared to control retina (0 days *ex vivo*), with reduced complexity (as assessed by Sholl analysis), total dendritic length, and dendritic field area. NAM treatment was highly effective, preserving dendritic morphology (**Figure 1J and K**). Dendritic varicosities were significantly increased in density as a function of dendrite length and dendritic field area, indicating dystrophy and degeneration of dendrites. NAM at a range of doses prevented these changes (**Figure 1M**). These data support previous findings on the protective effects of NAM, and extend its known effects to encompass neuroprotection across RGC compartments within the retina in additional animal models.

### Nicotinamide alters metabolic profiles in normal, non-diseased retinal ganglion cells

Given the importance of NAD in a wide array of metabolic processes, the potential of NAM as an interventional and prophylactic treatment in human glaucoma, we sought to determine whether NAM treatment alters RGC metabolic profiles under normal physiological conditions. We optimized a low molecular weight mass spectrometry platform to assess 73 essential low molecular weight metabolites across the RGC trajectory in normotensive rats (NT; normal IOP, no disease). We used bulk tissue (rat retina, optic nerve, superior colliculus (SC; the major RGC terminal in the rodent)) to avoid metabolic changes induced by cell sorting or tissue processing protocols. These data provide a novel metabolome across RGC compartments; dendrites, soma, and unmyelinated axons (retina), myelinated axons (optic nerve), and terminal arbors (SC). The normal rat metabolome varies significantly by these selected regions, with all 73 metabolites differentially abundant across tissues. Unsupervised hierarchical clustering (HC) separated samples into discrete groups that matched the different tissues, confirming globally distinct metabolic profiles (**Supplementary Figure 2A**). Metabolites were most highly abundant in the optic nerve, followed by the SC and retina respectively. Correlation of all individual metabolites for all samples demonstrated that the SC had the highest intra-tissue correlation (**Figure 2A**). In order to identify distinct metabolic signatures of the discrete tissues we used principle component analysis (PCA). Unsupervised PCA separated samples into discrete tissue groups supporting the HC data with PC1 describing the vast majority of variation (95.3%; **Figure 2B**). Creatine, hypotaurine, 2-aminobutyric acid, glycerophosphocholine and cysteine were the highest contributing loading factors on to PC1 demonstrating that relative abundance remained the largest determinant of tissue type even after scaling to reduce its effects (**Supplementary Figure 2B and C**). We specifically queried how NAD, NADH, and NAM vary across these tissues. NAD was higher in retina and optic nerve than in the SC, but NADH was much more abundant in the optic nerve. This results in a lower NAD:NADH ratio in the optic nerve. NAM was relatively low in the retina compared to both the optic nerve and SC (**Figure 2C**).

**Figure 2.**
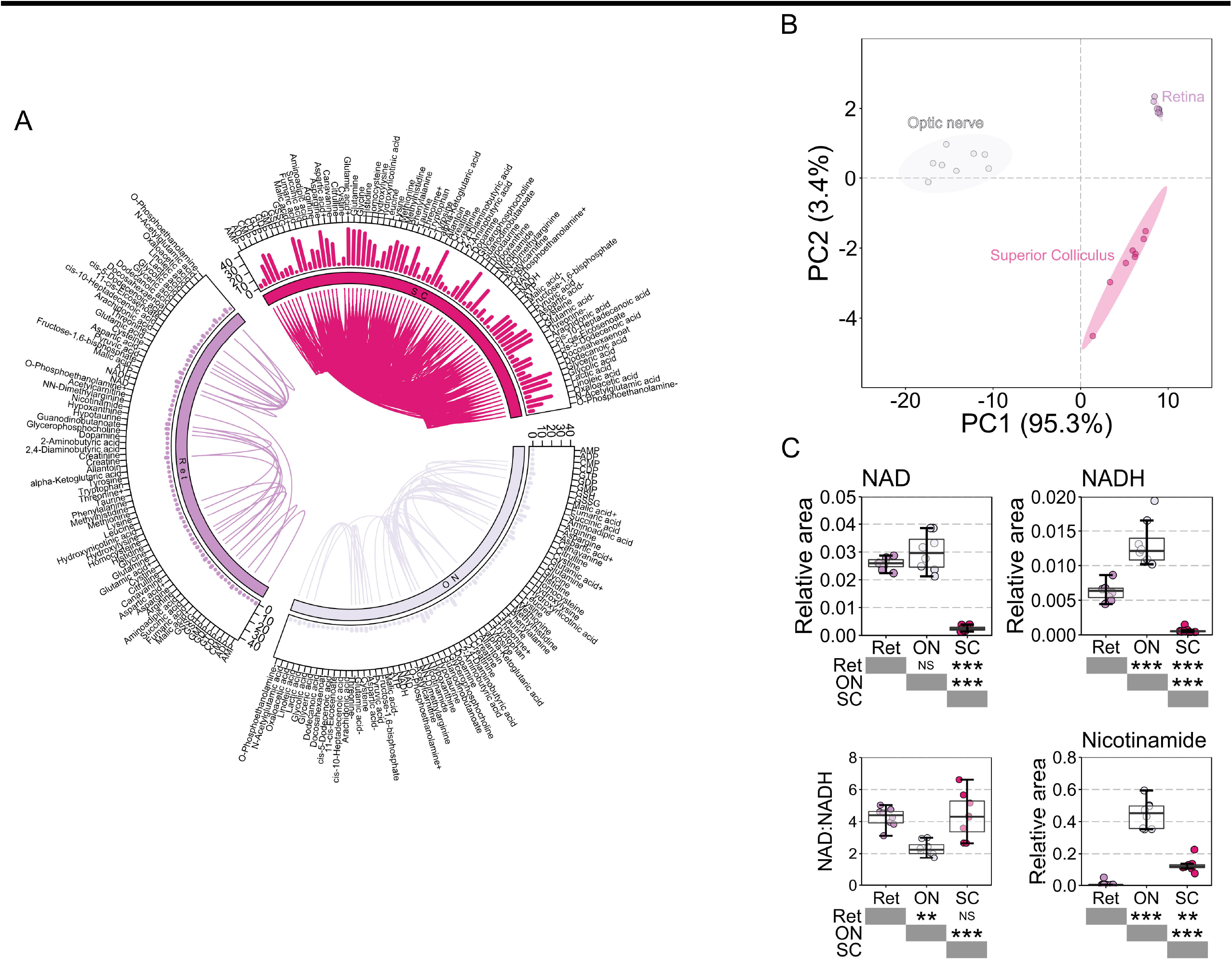
Metabolic profiles of retinal ganglion cell related tissues. (**A**) Seventy three low molecular weight metabolites could be reliably detected in retina (*n* = 8), optic nerve (*n* = 8) and superior colliculus (SC; *n* = 8 hemispheres) of Brown Norway rats. Within tissue correlation of metabolites is demonstrated by a circus plot with linkers between metabolites. The number of significantly correlated metabolites is shown as a bar graph in the outer circle. The SC (pink) showed the greatest degree of within tissue correlation of metabolites. (**B**) Tissue differences were explored by principle component analysis (PCA) which revealed a clear distinction of tissues predominantly along one component (PC1). (**C**) NAD and related metabolites NADH and nicotinamide were compared between tissues. The optic nerve was most abundant in NAD, NADH, and nicotinamide but had the lowest NAD:NADH ratio, whilst the retina was comparatively low in nicotinamide, and the SC comparatively low in NAD and NADH. ** *P* < 0.01, *** *P* < 0.001, NS *P* > 0.05.

NT rats treated with NAM 10 days prior to metabolomic analysis demonstrated a small but statistically significant reduced IOP of −1.2 mmHg (the physiological impact of which is likely to be limited) (**Figure 3A**). HC of individual samples demonstrated that NT-NAM treated retina were largely similar to normal NT retina, but with a clear distinction in metabolic profile in the optic nerve and SC (**Figure 3B**). PCA demonstrated that NAM induced changes were not sufficient to drive group distinction (**Figure 3B**). NAM treatment resulted in 9 changed metabolites (11%) in the retina, 24 in the optic nerve (30%), and 5 in the SC (6%) (**Figure 3C-F**). Increased NAD, NADH, and threonine were common to all tissues, with glyceric acid also increased in both retina and optic nerve (**Figure 3G and H**). These metabolites alone were sufficient to distinguish NT and NT-NAM samples by HC (**Figure 3G**). NAM was only increased in the optic nerve, suggesting that conversion to NAD may be saturated in the optic nerve at this dose (**Figure 3H**). The NAD:NADH ratio was only significantly altered in the optic nerve (40% increase), as NAD was increased to a greater proportion than NADH (~2.4 fold compared to ~1.7 fold), suggesting a larger pool of free NAD (**Figure 3H**). When cultured in NAM, dissociated neurons from retina, optic nerve, and cortex produce increased NAD over untreated cells demonstrating the ability to rapidly and directly metabolize NAM to NAD without the need for first pass metabolism (**Figure 3I**). Pathways analysis revealed that NAM induced changes are predicted to significantly impact nicotinate and nicotinamide metabolism across the retina and optic nerve. Arginine biosynthesis is predicted to be affected in the retina, and phenylalanine, tyrosine and tryptophan biosynthesis, taurine and hypotaurine metabolism, alanine, aspartate and glutamate metabolism, D-glutamine and D-glutamate metabolism, and glycine, serine and threonine metabolism are predicted to be altered in the optic nerve. In these pathways, significant impact was attributed predominantly to NAM and NAD or amino acids L-threonine, L-glutamate, L-aspartate, and phenylalanine. There were no predicted pathway changes in SC dataset (**Figure 3J**).

**Figure 3.**
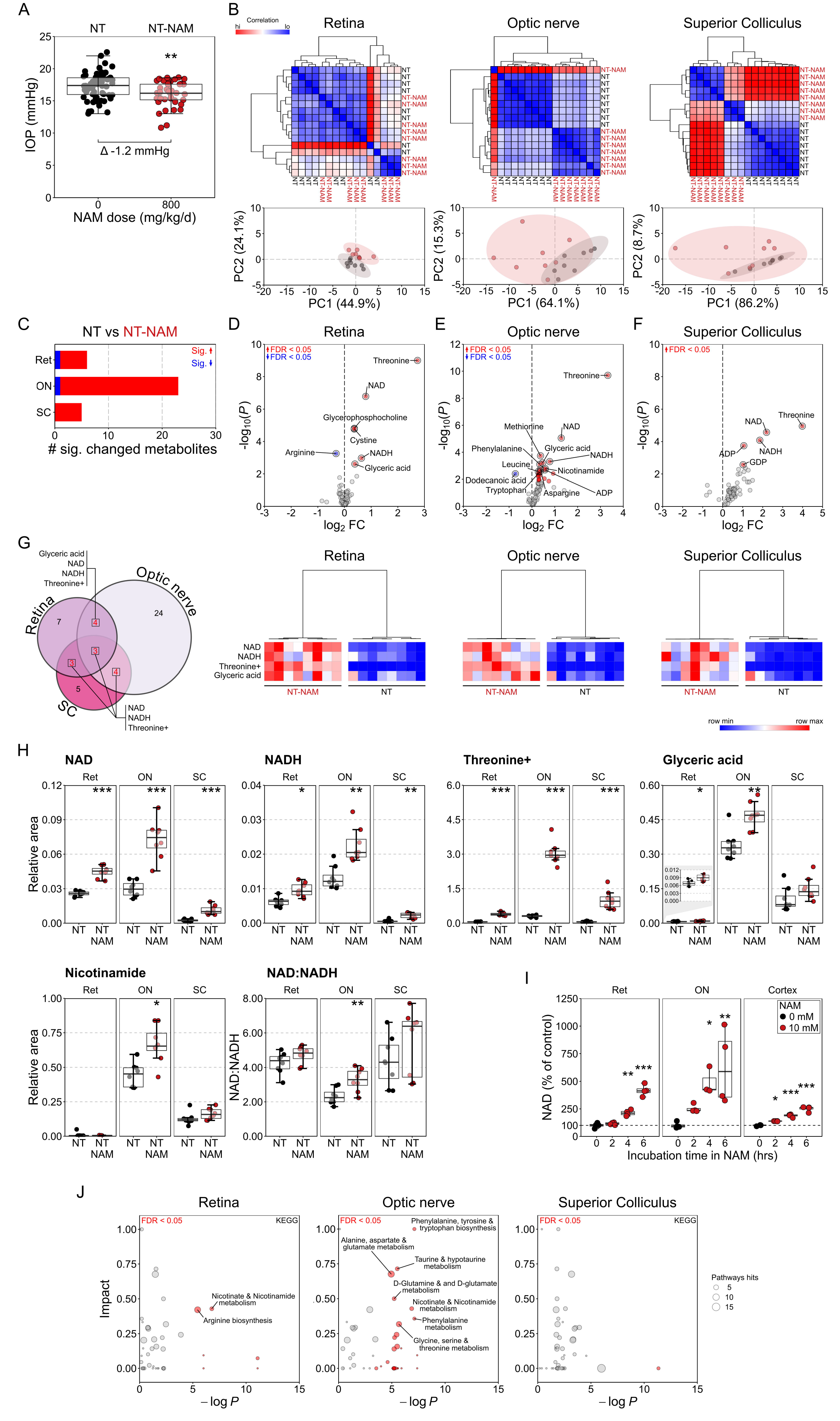
Nicotinamide alters metabolic profile of retinal ganglion cell related tissues. (**A**) High dose nicotinamide supplementation resulted in small significant decrease in IOP on average in NT eyes (*n* = 60 NT eyes, 40 NT-NAM (800 mg/kg/d) eyes). (**B**) BN rats were either untreated (NT) or maintained on NAM for 1 week (NT-NAM, 800 mg/kg/d) before metabolomics analysis of tissue. Unsupervised hierarchical clustering (HC) showed no clear distinction between NAM treated and untreated NT retina metabolic profiles (*n* = 8 NT retina, 8 NT-NAM), but clear separation in the optic nerve (*n* = 8 NT optic nerves, 8 NT-NAM) and superior colliculus (*n* = 8 NT SC hemispheres, 8 NT-NAM; all scaled where red = high correlation, blue = low correlation). Principle components analysis (PCA) showed little discriminating power indicating limited large scale NAM induced changes (NT = black, NT-NAM = red). (**C**) Comparison of NT and NT-NAM treated tissue revealed a number of significantly changed metabolites in **D** retina, **E** optic nerve, and **F**) superior colliculus (FDR < 0.05; red = increased in NT-NAM, blue = decreased), with the optic nerve the most changed. (**G**) Comparison of significantly changed metabolites across tissues revealed NAD, NADH, threonine, and glyceric acid (later in retina and optic nerve only) to be commonly increased (Euler plot showing total changed metabolites, with the number of common changed metabolites between tissues denoted in red). (**G**) HC dendograms and heatmaps (where red = highest value, blue = lowest value by row) and plotting of individual sample relative area (**H**) demonstrate the clear difference in these metabolites between NAM treated and untreated retina, optic nerve ad superior colliculus. (**H**) NAD was increased in all tissues, indicating that NAM dietary supplementation successfully translates to increased NAD in RGC relevant tissues. The optic nerve was the only tissue to show a change in the NAD:NADH ratio, indicting a potential saturation of conversion to NAD from NAM and a saturation of the NAD pool, or a greater recycling of NAD back to NAM. (**I**) Luminometry-based NAD assays on retina, optic nerve and cortex incubated with NAM showed that it can be converted to NAD within hours, increasing with increased exposure, suggesting an increase in the NAD pool (*n* = 4 retinal replicates where each is composed of 2 pooled retinas, for each time point a sample was taken from each replicate; *n* = 4 optic nerve replicates where 8 optic nerves were divided in to 1mm segments to give 16 independent samples for each timepoint; *n* = 4 cortex replicates where each replicates is a single cortex hemisphere and for each time point a sample was taken from each replicate). Optic nerve demonstrated the largest increase over control (untreated optic nerve) supporting the metabolomics observations. (**G**) Pathways analysis (KEGG) demonstrated that the changed metabolites (in **D-F**) resulted in minimal pathway changes of significant impact in the retina, and none in the superior colliculus. In the optic nerve, changes were predicted predominantly to NAD and related metabolite pathways. Pathways are highlighted red where FDR < 0.05, and annotated when in conjunction with high impact (*i.e*. predicted knock-on effects to the pathway); size denotes the number of metabolites within the pathway (scale adjacent). * *P* < 0.05, ** *P* < 0.01, *** *P* < 0.001.

### Nicotinamide protects against metabolic disruption following ocular hypertension

The retinal metabolome is significantly altered following 3 days of OHT, a time point at which IOP is high but no detectable RGC loss has occurred (15). HC revealed heterogeneity across and within conditions, but with a division between samples such that NAM treated eyes were largely grouped and distinguished (**Figure 4A**). In the retina OHT led to 25 changed metabolites relative to NT controls (24 increased, 1 decreased). Notable changes were increased alpha-ketoglutaric acid, homocysteine, threonine, NAD, and glycerophosphocholine, while creatinine was the sole reduced metabolite (**Figure 4B**). Comparison of OHT-NAM treated retinas to NT-NAM (thus excluding normal NAM induced changes) revealed that NAM treatment completely prevented any OHT induced metabolite changes (**Figure 4B**). NAD remained high in OHT-NAM retinas (**Supplementary Figure 3A**). Pathway analysis predicted changes to alanine, aspartate and glutamate metabolism, D-glutamine ad D-glutamate metabolism, arginine biosynthesis, glyoxylate and dicarboxylate metabolism, and histidine metabolism (**Figure 4C**).

**Figure 4.**
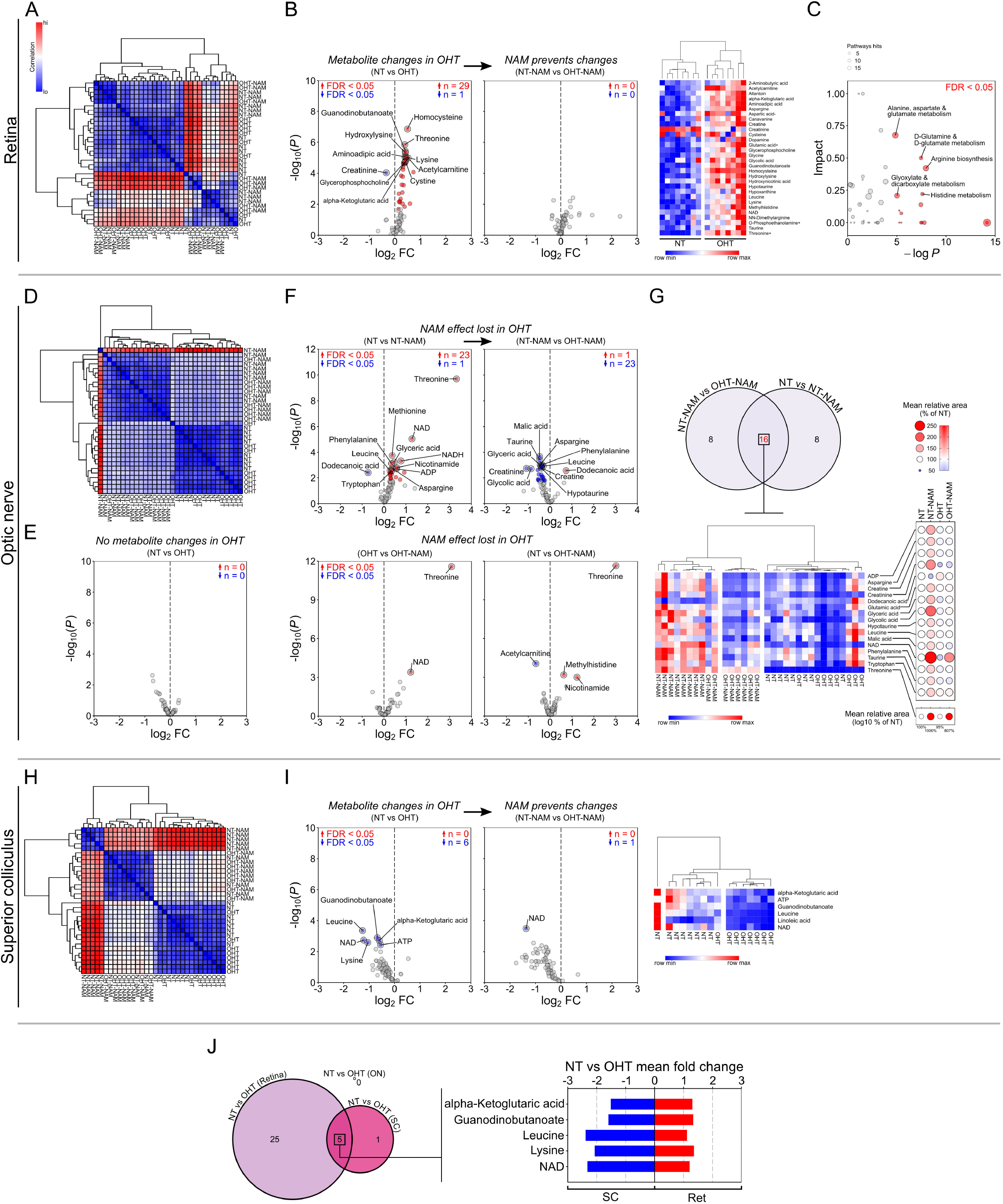
Nicotinamide prevents metabolic disruption. The metabolic profiles of BN rats following 3 days of OHT (prior to detectable neurodegeneration) were compared to NT controls, and the effects of pretreatment with NAM (800 mg/kg/d) explored (*n* = 8 retinas, optic nerves and superior colliculus hemispheres for each condition). (**A**) In the retina, unsupervised hierarchical clustering (HA) demonstrated that samples largely clustered based on exposure to NAM, but with heterogeneity across conditions (heatmap scaled where red = high correlation, blue = low correlation). (**B**) OHT induced a number of metabolic disturbances with a number of significantly increased metabolites (NT vs OHT; FDR < 0.05; red = increased in NT-NAM, blue = decreased; further shown in HA dendogram and heatmap; red = highest value, blue = lowest value by row); these changes were completely absent in NAM treated retinas (NT-NAM vs OHT-NAM; which controls for NAM specific effects). (**C**) Pathways analysis (KEGG) demonstrated a number of potentially effected pathways (pathways are highlighted red where FDR < 0.05, and annotated when in conjunction with high predicted impact, size denotes the number of metabolites within the pathway). (**D**) In the optic nerve, HA revealed clear distinction between NAM treated and untreated samples, irrespective of disease grouping. (**E**) No metabolites were significantly altered compared to NT following 3 days of OHT (NT vs OHT). (**F)** While NT-NAM treated optic nerves demonstrated a number of significantly changed metabolites over NT nerves, these effects were lost under OHT as demonstrated by the reversal of a number of these changes (NT-NAM vs OHT-NAM) and the similar profiles of OHT-NAM optic nerves to both OHT and NT nerves (OHT vs OHT-NAM, NT vs OHT-NAM respectively). (**G**) Comparison of NT-NAM vs OHT-NAM changes against NT vs NT-NAM changes, demonstrated 16 commonly changed metabolites (Euler plot showing total changed metabolites, with the number of common changed metabolites between comparisons denoted in red). HA of these metabolites separated individual NAM treated groups from controls, and plotting the mean abundance of these relative to NT control further highlights NAM specific changes that were reversed under OHT. (**H**) In the superior colliculus, HA largely distinguished individual groups, suggesting well defined metabolic profiles per condition. (**I**) In the SC, OHT resulted in a significant decrease of metabolites (NT vs OHT), which was prevented by NAM treatment (NT-NAM vs OHT-NAM). NAD in the SC was lower in OHT-NAM treated than NT-NAM treated (but remained above NT and OHT control levels; Supplementary Figure 3). (**J**) Comparison of OHT changes relative to NT in the retina, ON and SC revealed that 5 metabolites were commonly changed, where increases in the retina were mirrored by decreases in the superior colliculus (Euler plot showing total changed metabolites, with the number of common changed metabolites denoted; for mean fold change plot, red = increase, blue = decrease). * *P* < 0.05, ** *P* < 0.01, *** *P* < 0.001.

In the optic nerve, HC revealed discrete untreated and NAM groups (**Figure 4D**). No metabolites were significantly changed in OHT relative to NT (**Figure 4E**). In OHT-NAM, 24 metabolites were significantly lower than in NT-NAM. Comparison of OHT-NAM to untreated OHT and NT demonstrated only 2 and 4 changed metabolites respectively (**Figure 4F**). Collectively, these data suggest that the NAM induced metabolic profile change exhibited in normal tissue is lost during OHT. Comparison of NAM changes in NT (NT *vs*. NT-NAM) and OHT (NT-NAM *vs*. OHT-NAM) optic nerve demonstrated 16 commonly changed metabolites that reflect NAM specific changes (**Figure 4G**). HC using only these 16 metabolites was sufficient to separate individual NAM treated groups from untreated groups. Comparison of these changed metabolites relative to NT demonstrated marked increases in NT-NAM only, with OHT-NAM retuned to NT levels across all metabolites (with the exception of NAD and threonine) (**Figure 4G**). Crucially, NAD, NADH, and NAM remained higher in OHT-NAM optic nerves than in OHT, while NAD was reduced and NAM increased in OHT-NAM compared to NT-NAM (**Supplementary Figure 3B**).

In the SC, NAM treated conditions were clearly distinguished by HC (**Figure 4H**). During OHT 6 metabolites declined relative to NT controls, of which 5 of these changes were prevented in OHT-NAM (**Figure 4I**). NAM treatment in OHT was sufficient to replete NAD in the SC, with a diminished increase over normal NT levels relative to NT-NAM (**Supplementary Figure 3C**). The small number of metabolite changes in the SC were insufficient to drive any predicted pathways changes. Comparison of changes in OHT across tissues revealed 5 commonly changed metabolites in the retina and SC (**Figure 4J**). Of these, all were changed in opposing directions, demonstrating a concomitant increase in the retina and decrease in the SC. These data provide compelling evidence that retinal ganglion cell metabolism is altered early during glaucoma pathogenesis and can be prevented or buffered by oral NAM administration.

### Nicotinamide protects retinal ganglion cells from metabolic crisis

We have previously demonstrated dysregulated gene expression across multiple metabolic pathways (5), including widespread upregulation of genes encoding oxidative phosphorylation proteins (OXPHOS). Given that genetic changes coincide with mitochondrial cristae loss and increased cytochrome *c*, it can be hypothesized that these may reflect an upregulated transcriptional response in order to increase mitochondrial OXPHOS capacity. Gene expression in axotomized retina demonstrated a reduction in the mitochondrial-RNA:nuclear-RNA ratio suggesting a loss of mitochondrial capacity or a reduced number of mitochondria (**Figure 5A**). We queried whether the ratio of ATP:ADP was reduced in OHT, which would indicate a switch towards glycolysis from OXPHOS, how this may be affected by NAM treatment. No significant changes in this ratio were detected in the retina or optic nerve at 3 days in the rat OHT model. In the SC, the ATP:ADP ratio was significantly reduced in OHT and this was prevented by NAM treatment (**Figure 5B**). Since within condition variability was high, which may reflect mixed cell type-specific responses, we analyzed OXPHOS and glycolysis in purified mouse RGC cultures with NAM supplementation. Oxygen consumption rate was significantly increased in cells supplemented with 50 and 500 μM NAM, with the effect diminished at higher concentrations. No significant change in extracellular acidification rate was detected (**Figure 5C**). These findings suggest that NAM supplementation increases RGC OXPHOS capacity. We therefore tested the ability of NAM to protect RGCs against OXPHOS inhibition using an intravitreal rotenone model (inducing Complex I inhibition which results in the rapid degeneration of RGCs). Intravitreal injection *in vivo* of rotenone induced significant RGC death within 1 day, which was significantly reduced by NAM supplementation (**Figure 5D and E**). Thus, in the absence of axon degenerative stress, nicotinamide can provide a robust neuroprotection against mitochondrial stress.

**Figure 5.**
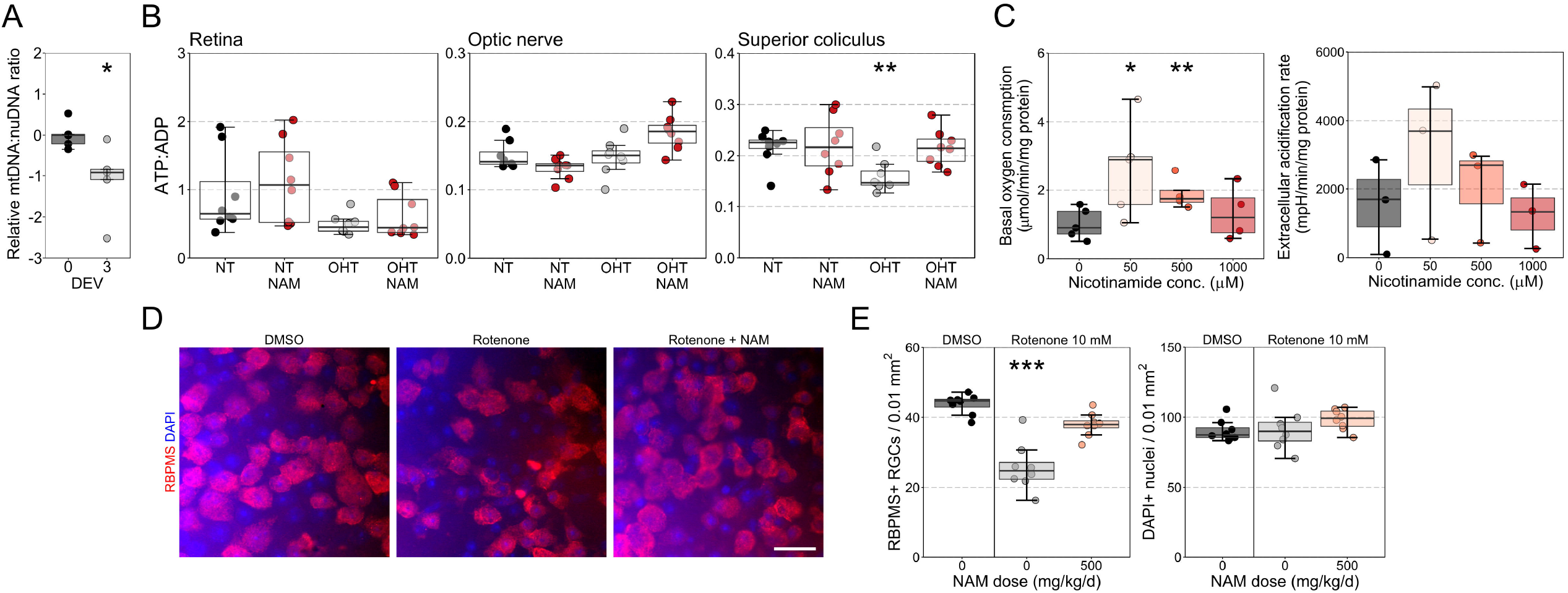
Nicotinamide buffers against metabolic crisis. (**A**) RGC axotomy resulted in a decrease in mitochondrial: nuclear derived DNA (mtDNA:nuDNA; relative expression of *mtCo2* and *rps18*) suggesting a loss of mitochondrial capacity or reduced numbers of mitochondria (*n* = five 0 days *ex vivo* (DEV) retinas, five 3 DEV). (**B**) The ratio of ATP:ADP was not significantly altered in the retina or optic nerve, but was reduced under OHT in the superior colliculus which could indicate a switch towards favoring glycolysis over oxidative phosphorylation (OXPHOS; *n* = 8 retina, optic nerve and superior colliculus hemispheres per condition). This was reversed under NAM treatment. (**C**) Oxygen consumption rate was significantly increased in cultured RGCs supplemented with 50 and 500 μM NAM, suggesting NAM increases OXPHOS capacity (*n* = 5 cultures with 0 μM NAM, 5 cultures with 50 μM NAM, 4 cultures with 500 μM NAM, and 4 cultures with 1000 μM NAM). Extracellular acidification rate did not change significantly (*n* = 3 cultures per condition). (**D**) Intravitreal injection of Rotenone (causing rapid depletion of OXPHOS) caused significant loss of RGCs after 24 hours, and pre-treatment with NAM provided significant protection against this loss (*n* = 8 retina per condition). Scale bar in D = 20 μM. * *P* < 0.05, ** *P* < 0.01, *** *P* < 0.001.

### Nicotinamide increases mitochondrial size and motility

Mitochondrial remodeling and/or loss is a common feature of neurodegenerative disease which may drive neuronal metabolic dysfunction. We directly assessed changes to mitochondrial morphological integrity after 14 days of OHT in the rat. TOMM20 labelled mitochondria were imaged by high resolution Airy confocal microscopy (35 nm / px) and reconstructed (**Figure 6A**). Mitochondrial volume and surface area were significantly reduced in the ganglion cell layer (GCL) and RNFL (comprising RGC soma and axons and other retinal neurons/glia) of OHT animals (**Figure 6B**). NAM treatment partially protected from these mitochondrial degenerative changes. In OHT animals, mitochondrial volume and surface area were significantly increased in the IPL (comprising RGC dendrites and interneuron neuropil), and further increased in OHT-NAM treated animals (**Figure 6C**). These different changes between retinal layers may be representative of the various cell types that make up these layers, their neuronal networks, and how they respond to OHT. To determine RGC specific mitochondrial responses to injury and NAM, we utilized a novel mitochondrial reporter mouse where YFP is expressed under a rat *Eno2* promoter (neuron specific) and localized to mitochondria via a *Cox8a* gene-targeting signal fused to the YFP N-terminus (neuronal expression shown in **Supplementary Figure 4**). In this strain (designated MitoV; for visual tissue), YFP expression is RGC specific in the inner retina, allowing the assessment of mitochondria specifically in RGCs (with expression in a subset of bipolar cells and photoreceptor outer segments which can be optically dissected; **Supplementary Figure 4A-C**). We hypothesized that physical axonal damage and mitochondrial stress work in concert to drive vision loss in glaucoma. To assess each of these in an isolated context, we induced RGC axotomy (uncoupling the RGC from the terminal visual thalami) or acute, severe mitochondrial dysfunction (intravitreal rotenone injection). Using the retinal explant model to induce an RGC specific injury, we assessed mitochondrial morphology in the GCL/RNFL (**Figure 6D**). At 12 hours *ex vivo* mitochondrial volume and surface area were significantly increased. NAM supplementation of the media exaggerated the increase in mitochondrial volume and surface area **(Figure 6E**). Intravitreal injection of rotenone resulted in a significant reduction of mitochondrial volume and surface area in the IPL at 1 day post injection. NAM supplementation was unable to prevent these morphological changes even though NAM treatment was grossly neuroprotective (**Figure 6F and G**). These data suggest that even across a range of insults, mitochondria increase or maintain their size when NAM is present. However, these changes may reflect the tissue culture conditions (*e.g*. Ca^2+^ buffering deficits or deviation from normoxia in the axotomy model) or the abrupt complete physiological loss of mitochondrial Complex I function (in the rotenone model) which are exaggerations of the intrinsic RGC injury responses during glaucoma pathogenesis.

**Figure 6.**
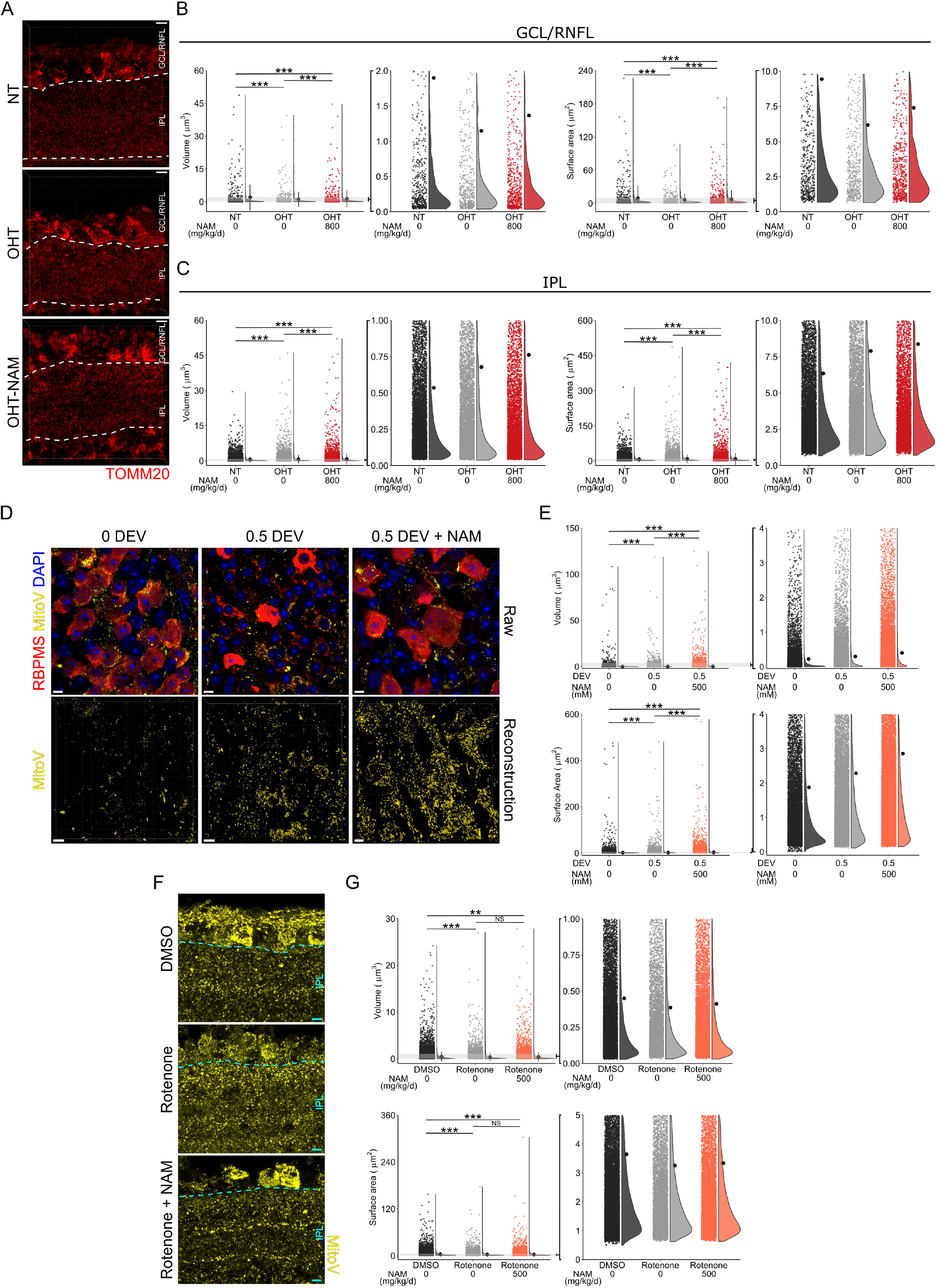
Nicotinamide increases mitochondrial size. (**A**) Rat retina following 14 days of OHT (untreated (OHT, *n* = 6 retina) and NAM treated (800 mg/kg/d, OHT-NAM, *n* = 4 retina)) and NT control (*n* = 4 retina) tissue was cryo-sectioned, mitochondria labelled by anti-TOMM20, and imaged by Airy confocal scanning microscopy. The ganglion cell layer/retinal nerve fiber layer (GCL/RNFL) and inner plexiform layer (IPL) were analyzed separately (boundaries demarcated by white broken lines). (**B**) Mitochondrial volume and surface area were significantly reduced under OHT in the GCL/RNFL and this was mitigated by NAM (*n* = 535 disconnected volumes in NT, 490 in OHT, 652 in OHT-NAM). (**C**) In the IPL mitochondrial volume and surface area increased under OHT, which was exaggerated under NAM treatment (*n* = 9729 disconnected volumes in NT, 8026 in OHT, 6413 in OHT-NAM). (**D**) Retinal axotomy culture of MitoV mice (RGC specific YFP labeled mitochondria in the inner retina) maintained for 0.5 days *ex vivo* (DEV) with or without NAM treated media (and controls fixed at point of euthanasia, 0 DEV) allowed imaging of RGC specific mitochondria in the GFL/RNFL by Airy confocal scanning microscopy following flat mounting (*n* = 4 retina for all conditions). (**E**) The mean mitochondrial volume and surface area were increased under axotomy, with a further increase in size under NAM treatment (*n* = 15216 disconnected volumes in 0 DEV, 16765 in 0.5 DEV, 24558 in 0.5 DEV + NAM). (**F**) The effects NAM at mitigating direct mitochondrial stress were investigated by intravitreal rotenone injection in MitoV mice (*n* = 9 DMSO vehicle treated retina, 3 Rotenone, 7 Rotenone + NAM (500 mg/kg/d)). Tissue was cryo-sectioned and imaged by Airy confocal scanning microscopy and mitochondria within the IPL (boundary demarcated by cyan broken line). (**G**) Rotenone caused a reduction in mitochondrial volume relative to controls (DMSO vehicle) in the IPL. NAM had little effect is preventing this change (*n* = 15380 disconnected volumes in DMSO vehicle treated retinas, 7987 in rotenone treated, 10418 in rotenone + NAM). These results demonstrate the heterogeneous nature of OHT insult on mitochondria that shares features with both direct RGC and direct mitochondrial stress. Scale bar in A = 5 μm in A, D, and F. * *P* < 0.05, ** *P* < 0.01, *** *P* < 0.001.

In order to differentiate NAM-induced and insult-induced mitochondrial changes we administered NAM in the diet of uninjured control MitoV mice for 1 week prior to fixation and imaging (**Figure 7A**). RGC mitochondria in the GCL/RNFL increased in volume and surface area compared to untreated controls, suggesting that the increase in volume and surface area observed in the NAM treated injury paradigms were NAM specific responses (**Figure 7B**). The presence of larger mitochondria prior to damage may therefore explain the maintenance of mitochondrial volume and surface area which was observed in the rat GCL/RNFL in NAM treated injuries and the exaggerated mitochondrial expansion seen in the rat IPL. *In vivo* imaging of MitoTracker labeled mitochondria in purified cultured RGCs (**Figure 7C**) demonstrated an increase in mitochondrial size following NAM supplementation (24 hours prior) which was dose dependent (**Figure 7D**). Mitochondria occupied a greater proportion of neurite length (**Figure 7D**). Live imaging, analyzed as kymographs (**Figure 7E**), revealed a greater number of mitochondria and a striking increase in mitochondrial population mobility and velocity across NAM doses over untreated control (**Figure 7F**). Total and stationary mitochondria displayed some variability with NAM dose (**Figure 7F**). Larger, more mobile mitochondria may be indicative of increased mitochondrial fusion, with implications for increased OXPHOS capacity (as supported by the increased oxygen consumption rate measured by Seahorse analysis) and maintenance of energy demands across the cell.

**Figure 7.**
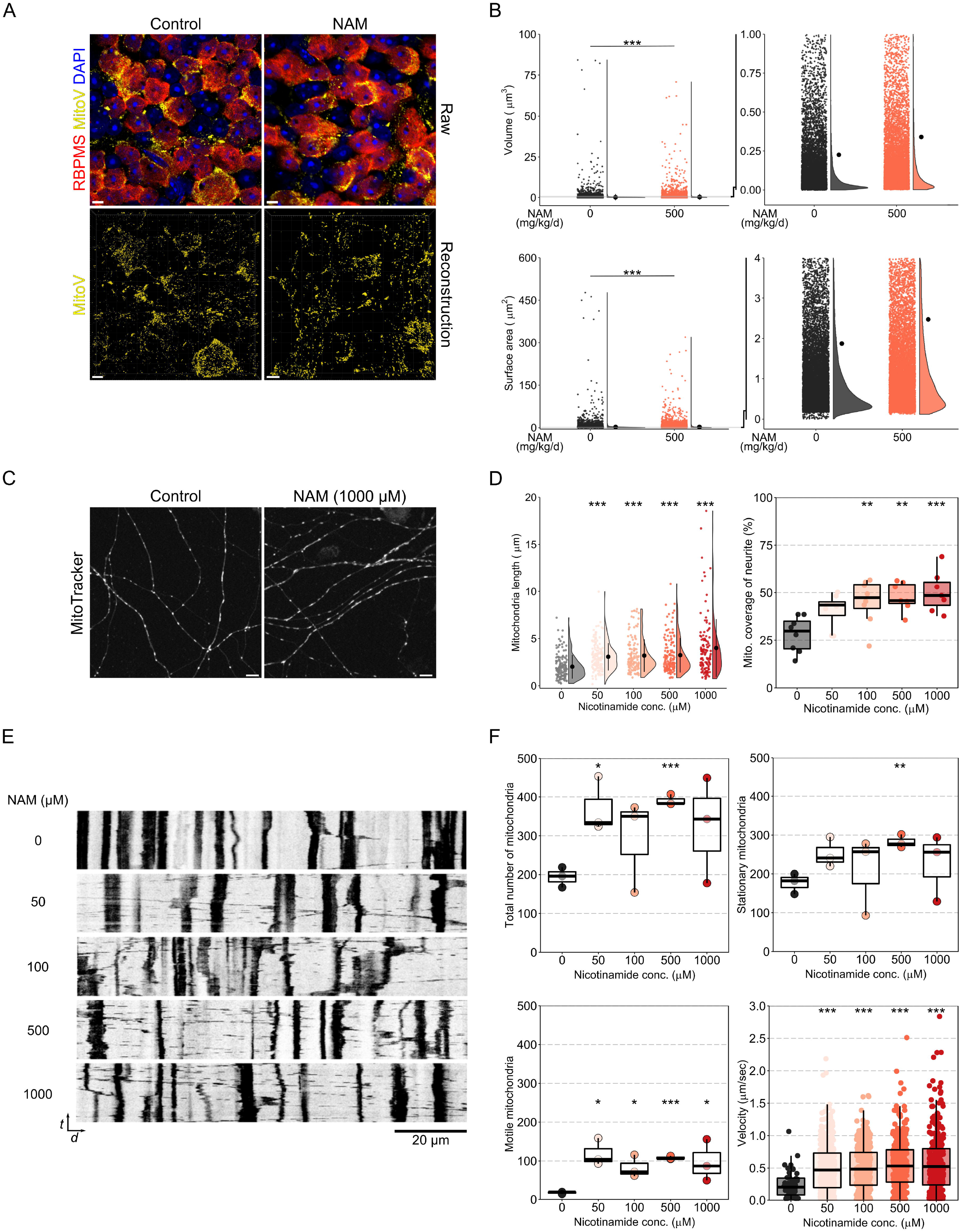
Nicotinamide increase mitochondrial size and motility in normal RGCs. (**A**) In order to determine whether NAM increases mitochondrial size in uninjured retina, MitoV mice were maintained on a NAM diet (500 mg/kg/d) for 1 week. Retina were fixed immediately after euthanasia and compared to untreated controls by Airy confocal imaging and reconstruction (*n* = 4 control retinas, 4 NAM supplemented). (**B**) NAM supplementation increased mean mitochondrial volume and surface area in RGCs, supporting observations of mitochondria under stress above (*n* = 15216 disconnected volumes in control, 13306 in NAM supplemented). (**C**) In order to observe these NAM effects on mitochondria *in vivo*, RGC cultures were grown and exposed to NAM treated media for 24 hours. MitoTracker labelled mitochondria were time-lapse imaged over 10 minutes (example images shown in **C** and as kymographs in **E** where the y axis = time (*t*) and the x axis = distance long the neurite (*d*)). Morphology was examined in the first frame and demonstrated that mitochondrial length was increased with increasing NAM concentration (**D**; n = 148 mitochondria in control, 154 in 50 μM NAM, 141 in 100 μM NAM, 140 in 500 μM NAM, 122 in 1000 μM NAM; mitochondria from 8 neurites from 3 independent cultures for all conditions). Mitochondria occupied a greater percentage of neurite length in NAM exposed retina (**D**; *n* = 8 neurites from 3 independent cultures for all conditions), which was partially explained by an increase in observed mitochondria (**F**). Analysis of mitochondrial movements demonstrated a greater number of both mobile and stationary mitochondria (**F**; *n* = 3 independent cultures for all conditions). Mitochondrial velocity was significantly increased with NAM supplementation (**F**; n = 52 mitochondria in control, 356 in 50 μM NAM, 301 in 100 μM NAM, 324 in 500 μM NAM, 292 in 1000 μM NAM; mitochondria from 3 independent cultures for all conditions). Since a greater number of stationary mitochondria were observed, this suggests that the increase in mitochondria numbers may be a factor of increase mitochondrial biogenesis. Scale bar = 5 μm in A, 10 μm in C and 20 μm in E. * *P* < 0.05, ** *P* < 0.01, *** *P* < 0.001.

### Nicotinamide reduces retinal ganglion cell metabolic demands

Since OXPHOS capacity and mitochondrial motility increased following NAM application, we questioned whether RGC electrical activity was altered. Mouse retina were explanted and incubated with NAM for 2 hours before patch clamping (example profiles displayed in **Figure 8A**). At low NAM doses (1-10 mM) the action potential (AP) firing rate was slightly increased, with persistent firing at higher current inputs. Higher NAM doses (100-250 mM), in which we demonstrated neuroprotection in explant retina, demonstrated marked reduction in AP firing rates across current inputs (**Figure 8B**). The firing rate (I/O relationship AUC), AP amplitude, and AP rise time were significantly reduced at higher NAM doses (**Figure 8C**). Resting membrane potential, AP threshold, and rheobase were not affected by NAM suggesting integrity of the cell’s capacity to depolarize (**Figure 8D**). These data suggest that the higher neuroprotective doses of NAM lower AP firing frequency in RGCs, without compromising their potential to fire, and thus may reduce RGC metabolic demands.

**Figure 8.**
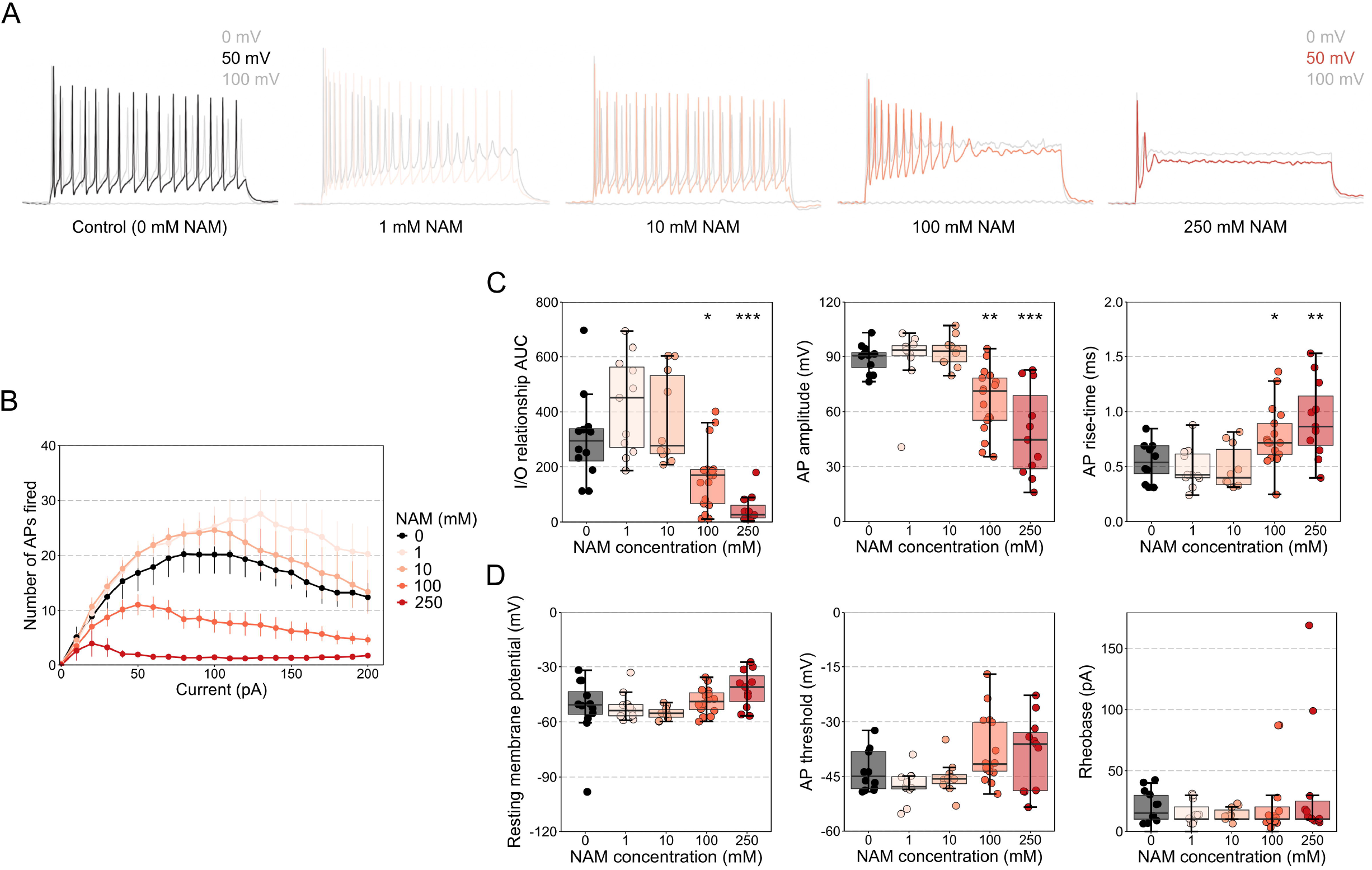
Nicotinamide lowers action potential firing frequency in RGCs. (**A**) Mouse retina were explanted and incubated with NAM for 2hrs before patch clamping. Example traces for individual RGCs are shown for each treatment group (0 mV profile in grey, 50 mV profile in color, 100 mV profile in grey; total RGCs = twelve control, eleven 1 mM NAM, ten 10 mM NAM, 17 100 mM NAM, eleven 250 mM NAM, from 3 retinas for each condition). (**B**) Action potential (AP) firing frequency was markedly reduced at higher doses of NAM (analogous to those that provide neuroprotection in the axotomy explant model). (**C**) The relationship of input current to output demonstrates a slight increase in AP firing at high current for low dose NAM (1 and 10 mM) with a significant reduction in firing across a large range of input currents for higher doses (100 and 250 mM). (**C**) AP amplitude was significantly reduced at the higher doses and AP risetime was significantly increased. (**D**) NAM did not affect the resting membrane potential, threshold for achieving an AP or the Rheobase indicating that the potential/ability to fire was not altered. Reducing AP firing may lower the metabolic demands of RGCs thus contributing to its neuroprotective effects. * *P* < 0.05, ** *P* < 0.01, *** *P* < 0.001.

## Discussion

The control of metabolic homeostasis is essential for normal physiology as well as during stress- and disease-related episodes. To assess this in the context of ocular disease, we introduce a novel, small molecular weight metabolome for the retina, optic nerve, and superior colliculus of the Brown Norway rat (*Rattus norwegicus*; the genetic standard for rat). We demonstrate that ocular hypertension causes early metabolic dysregulation that is prevented by nicotinamide treatment. These data advance the understanding of nicotinamide as a treatment for glaucoma, by demonstrating substantial metabolic protection and robust mitigation of glaucoma-related insults across multiple retinal ganglion cell compartments. Elevated IOP is a major risk factor for glaucoma and is the most significant factor in many patients. We modelled glaucoma in this context with an acute, inducible IOP increase (ocular hypertension) which drives long-term retinal and optic nerve NAD decline in the absence of age-related effects on NAD homeostasis. Nicotinamide treatment robustly protected against retinal ganglion cell damaging events. Metabolomic analysis confirmed that retina, optic nerve, and superior colliculus all demonstrated increased NAD following nicotinamide administration. Importantly, nicotinamide drove increases in NAD in mature *in vivo* systems and in isolated neurons demonstrating that retinal neurons themselves are capable of utilizing nicotinamide without a requirement for first-pass metabolism or other structural chemical changes. Crucially, high dose nicotinamide caused only minor changes to the normal metabolome, mostly influencing metabolites linked directly to NAD or increasing amino acids L-threonine, L-glutamate, L-aspartate and phenylalanine with limited predicted pathway effects (i.e. low predicted impact on critical pathway nodes via KEGG analysis), suggesting that chronic nicotinamide treatment is unlikely to have substantial deleterious effects in visual system tissue metabolism long-term. This is important if nicotinamide is to be considered as a prophylactic in glaucoma patients where significant neurodegeneration is yet to occur. Hierarchical clustering revealed that tissues could be grouped based on the presence or absence of nicotinamide treatment over disease related changes. However, this was largely driven by large fold changes in a small number of metabolites in nicotinamide treatment as opposed to smaller fold changes in numerous metabolites in ocular hypertension. Metabolic stabilization and neuroprotection were achieved with relatively short-term administration of nicotinamide (1 week prophylactic), and dissociated neurons were capable of mobilizing nicotinamide to NAD rapidly (within hours), demonstrating the short period required to achieve an increase in cellular NAD pools. These results support nicotinamide supplementation as a therapeutic strategy for neuroprotection in glaucoma with particular regard to its rapid mobilization to NAD in visual system tissues.

The maintenance of cellular NAD pools is critical for neuronal survival as it provides co-enzyme resources necessary for maintaining a variety of complex cellular functions including ATP production, protein deacetylase activity, calcium homeostasis, conversion to NADPH, and the maintenance of gene expression and DNA repair (16, 17). The presence of metabolic dysregulation and ATP depletion (5, 18), calcium dysregulation (19), mitochondrial degeneration (4, 5), and reactive oxygen species (20) all contribute to glaucoma pathogenesis, supporting a role for NAD as a critical regulator of neuronal health in glaucoma. Nicotinamide supplementation is, therefore, likely to confer multiple neuroprotective effects. These features are not unique to glaucoma and are common to other neurodegenerative diseases including mitochondrial degeneration and metabolic dysfunction. Increasing NAD through multiple substrate avenues has protective effects in many of these diseases, including Alzheimer’s disease (21–24), Parkinson’s disease (25–28), other neurodegenerations (29, 30), and other mitochondrial degenerative disorders (31, 32). Continued elucidation of the mechanisms of Wallerian degeneration has further demonstrated the role of NAD in axon survival. NAD decreases in damaged axons (33), while expression of the *Wld^S^* allele or viral overexpression/modified localization of NMNATs are protective, restoring NAD levels (5, 34–37). Increased NAD appears to exhibit numerous functional local roles in this context, including providing sufficient substrate to mitigate local SARM1-dependent NAD consumption and protecting against nicotinamide mononucleotide (NMN) induced axon damage (38, 39).

The extent to which the NAD increase was restricted to retinal ganglion cells is not known since the metabolomics data is derived from a mixed cell population which included retinal ganglion cell compartments. The optic nerve, containing retinal ganglion cell axons, represents the most retinal ganglion cell enriched tissue based on the relative abundance of cell types, suggesting that retinal ganglion cells likely exhibit increased NAD following nicotinamide supplementation. Increasing NAD across multiple cell types, including glia which offer metabolic support to retinal ganglion cells, may enhance the neuroprotective effect, yet this remains to be explored and is an exciting avenue for therapeutic research, especially in other neurodegenerative diseases in which compromised metabolic function is coupled with a neuroinflammatory components such as Alzheimer’s disease. Increasing NAD production by *Nmnat1* gene therapy in retinal ganglion cells is neuroprotective and supports a hypothesis in which retinal ganglion cell specific strategies to increase NAD are neuroprotective (5). Since the intracellular location of NAD is key to its neuroprotective effects (40) further studies examining this in retinal ganglion cells will aid in elucidating the mechanisms of neuroprotection. Approaches such as computational molecular phenotyping using NAD specific probes or live cell FRET imaging will further this understanding.

We identify mitochondrial protection and oxidative phosphorylation maintenance as potential routes to buffering retinal ganglion cells bioenergetic insufficiency (41–43). A concomitant decrease in retinal ganglion cell metabolic demand through reduced action potential firing frequency could mitigate the effects of a retinal ganglion cell energy crisis. A reduced energetic load on retinal ganglion cells may partially explain enhanced survival. Exogenous nicotinamide and NAD have demonstrated similar suppression and prolongation of action potential firing and activity across neuronal and cardiac tissue preparations (44–48). It is important to note that nicotinamide has been used therapeutically at high doses long-term in clinical preparations without noted incidences of visual or cognitive disturbances (9, 43). Pattern electroretinogram (ERG) is a retinal ganglion cell specific functional test, representing a global summation of retinal ganglion cell signals. Nicotinamide supplementation was sufficient to preserve pattern ERG under chronic glaucomatous pathology (5, 49) and has demonstrated a recovery in visual function as assessed by ERG in human glaucoma patients (7). These findings suggest that even if the changes in retinal ganglion cell activity identified by patch clamping were recapitulated *in vivo*, visual function would not be disturbed. While the rodent and human visual system share many common features, and NAD synthetic pathways are conserved, the metabolic consequences in humans could vary. As we focused on small molecular weight metabolites, the consequences to the whole metabolome remain unknown and further metabolomics and lipidomic studies in humans will be necessary especially within the context of clinical trials and long-term testing. To our knowledge, this is the first small molecular weight metabolome for these tissues which will provide a resource which may aid others in the identification of key metabolic changes related to visual pathologies.

These findings advocate furthering clinical trials for the use of nicotinamide as a treatment for glaucoma. Nicotinamide has a long established clinical safety profile, with limited long-term side effects. Long-term clinical trials that include individuals with early glaucoma (in which cell loss is heterogeneous and a significant portion of healthy cells likely remain) will be essential in determining the potential for nictoinamide as a prophylactic treatment for glaucoma. However, longer-term treatment of nicotinamide in animals retain their importance, especially in non-human primates which closely recapitulate the human visual system. A number of disease preventative measures have been identified in animal models of glaucoma, which provide robust treatment at early and mid-disease time points, but long-term neuroprotective effects are not maintained or are yet to be demonstrated (50–54). Animals were treated with high doses of nicotinamide that represent achievable scaled doses to humans (the lowest neuroprotective dose here of 200 mg/kg/d in the rat representing 1.9 g/day in a 60 kg human (11)). While we did not test any extra-visual system effects of these doses, the safety of the doses suggested here, as well as doses up to 12 g/d, is already established (9). Although some patients may find adherence to these doses difficult to achieve because of the necessary number of tablets (especially when taken in addition to other common medications typical of more elderly patients) and the regularity of dosing, doses in the 1-3 g/d range are likely to be suitable for many patients. Supporting this, 3 g/d (6 x 500 mg capsules) had a high tolerance and compliance in Hui *et al*. nicotinamide glaucoma clinical trial (94% adherence).

Collectively, our data suggest that nicotinamide given as an early intervention or prophylactic treatment produces minimal short-term changes to healthy, non-disease visual system tissue with no detectable deleterious effects to normal retinal ganglion cells. Nicotinamide buffers against metabolic and bioenergetic insufficiency to provide a potent neuroprotection against a variety of glaucoma-related stresses. The doses used here are realistic for human treatment regimens and support further clinical trials for glaucoma as well as potentially a range of metabolic and ophthalmic diseases.

## Materials and Methods

### Animal strain and husbandry

All breeding and experimental procedures were undertaken in accordance with the Association for Research for Vision and Ophthalmology Statement for the Use of Animals in Ophthalmic and Research. Individual study protocols were approved by Stockholm’s Committee for Ethical Animal Research (10389-2018). Animals were housed and fed in a 12 h light / 12 h dark cycle with food and water available *ad libitum*. Adult, male Brown Norway rats (BN, aged 12-20 weeks, weighing 300-400 g, SCANBUR) were utilized for inducing ocular hypertension (detailed below). C57BL/6J and MitoV (strain details and generation detailed below) mouse strains were bread and utilized at 12-20 weeks of age. For rats, Nicotinamide (NAM, PanReac AppliChem) was dissolved in drinking water (200 mg/kg/d, based on average consumption). Higher doses were achieved by combining treated drinking water and a custom diet (3130 ppm NAM to achieve 400 mg/kg/d, or 9380 ppm NAM to achieve 800 mg/kg/d based on a pre-determined g/d consumption of chow, base RM3 P diet, SDS). For mice, NAM was dissolved in drinking water (500 mg/kg/d based on average consumption). Water was protected from light and changed every 3-5 days and chow was maintained as necessary; both were provided in sufficient quantity to be available *ad libitum*.

### Induction and monitoring of rat ocular hypertension

Rat ocular hypertension was induced using a paramagnetic bead model as previously described (15). Rats were anesthetized with an intraperitoneal injection of ketamine (37.5 mg/kg) and medetomidine hydrochloride (1.25 mg/kg). Microbeads (Dynabead Epoxy M-450, Thermo Fisher) were prepared in 1x Hank’s balanced salt solution (HBSS −CaCl_2_ −MgCl_2_ −phenol red, Gibco) and 6-8 μl of bead solution was injected into the anterior chamber. Beads were directed with a magnet to block the iridocorneal angle. Rats received either bilateral injections (OHT) or remained bilateral un-operated (naïve), normotensive controls (NT). For rats pre-treated with NAM, NAM was given 1 week prior OHT induction (at either 200, 400, or 800 mg/kg/d) and maintained throughout the duration of the experiment. Intraocular pressure (IOP) was measured using a rebound tonometer (Tonolab, Icare). Rats were habituated to tonometry in the week prior to induction. Baseline IOP was recorded the morning before surgery and every 2-4 days afterwards. Rats were awake and unrestrained for tonometry. IOP recordings were always performed between 9 and 10am to avoid the effects of the circadian rhythm on IOP.

### OCT imaging

Optical coherence tomography (OCT) was performed at 14 days post IOP in BN rats, prior to euthanasia (n = 9 eyes NT, 7 OHT, 8 OHT treated with 200 mg/kg/d NAM (OHT-NAM)). Pupils were dilated with topical Phenylephrine Hydrochloride (2.5% w/v) eye drops before anesthesia was induced with an intraperitoneal injection of ketamine and medetomidine hydrochloride as above. OCT imaging was performed using a Phoenix Micron IV (Phoenix) with image-guided OCT attachment. B-scans centered on the ONH were captured in the horizontal and vertical plane. Image analysis was performed in MATLAB (MATrix LABoratory). First the retinal pigment epithelium and internal limiting membrane were delineated manually. In the presence of cupping, the cup depth was measured automatically as the vertical distance between the cup base and midpoint of the neuroretinal peaks. This measurement was used to determine loss of neuroretinal rim.

### Retina axotomy explant model

A retinal axotomy model was established with modification form previously described protocols (13, 14). Mice were euthanized by cervical dislocation and eyes immediately enucleated. Retinas were dissected free in cold HBSS and flat mounted on cell culture inserts (Millicell 0.4 μm pore; Merck) ganglion cell layer up. Retinas were maintained in culture (37°C, 5% CO_2_) fed by Neurobasal-A media supplemented with 2 mM L-glutamate (GlutaMAX, Gibco) 2 % B27, 1 % N2, and 1% penicillin/streptomycin (all Gibco) in 6-well culture plates. Half the media volume was changed after 2 days. For NAM treated retinas, NAM was dissolved in the culture media to a concentration of 100mM or 500mM. Retinas were removed from culture for further processing after 12 hours, 1 day, 3 days or 5 days *ex vivo* (DEV).

### Intravitreal rotenone model

Rotenone toxicity induced degeneration was induced in the retina following established protocols (55, 56). Mice were anesthetized with an intraperitoneal injection of ketamine (37.5 mg/kg) and medetomidine hydrochloride (1.25 mg/kg). Bilateral intravitreal injections were performed using a XG tri-beveled needle on a 10 μl glass syringe (WPI). One μl of a 10 mM rotenone (MP Biochemicals) solution dissolved in DMSO (PanReac AppliChem) or DMSO only (vehicle only control) was injected into the vitreous and the needle maintained in place for 30 seconds for distribution. For NAM pre-treated retinas, NAM (500 mg/kg/d) was administered in drinking water 1 week prior to rotenone injection. Mice were euthanized 24 hrs after injection by cervical dislocation and eyes immediately enucleated for further processing.

### Cryo-sectioning

Rats were heavily anaesthetized by intraperitoneal injection of pentobarbital (75 mg/kg), and euthanized by cervical dislocation. Mice were euthanized by cervical dislocation. Eyes for cryosectioning were enucleated, and fixed by immersion in 4.7% PFA for 24 hrs. The tissue was cryoprotected by immersion in 30% sucrose for 24 hrs before freezing in optimal cutting temperature medium (Sakura) on dry ice. Blocks were maintained at −80°C until use. 20 μm cryo-sections were cut (anterior to dorsal plane) using a cryostat (Cryostar NX70, Thermo Scientific) and stored at - 20 °C.

### Immunofluorescent labelling

Immunofluorescent labelling followed a standard protocol for cryo-sections and flat mount retina with the following exceptions. Cryo-sections were air dried for 15 min and rehydrated in 1M PBS for 5 min before following the protocol. Antibodies are detailed in **Supplementary Table 4**. For the standard protocol, tissue was isolated using a hydrophobic barrier pen (VWR), permeabilized with 0.5% Triton X-100 (VWR) in 1M PBS for 30 mins, blocked in 5% bovine serum albumin (Fisher Scientific) in 1M PBS for 30 mins, and primary antibody applied and maintained overnight at 4 °C. 5 x 5 min washes in 1M PBS were performed before applying secondary antibodies and maintaining at room temperature for 2 hours. Tissue was washed as before and DAPI nuclear stain (1 μg/ml in 1M PBS) was applied for 10 mins. Tissue was then washed once in PBS before mounted using Fluoromount-G and glass coverslips (Invitrogen). Slides were sealed with nail-varnish.

### Analysis of RGC degeneration

RGC loss and nuclear shrinkage was assessed through RBPMS and DAPI labelling of flat mounts. Images were acquired on a Zeiss Axioskop 2 plus epifluorescence microscope (Karl Zeiss). Six images per retina (40X magnification, 0.25 μm/pixel) were taken equidistant at 0, 2, 4, 6, 8, and 10 o’clock about a superior to inferior line through the optic nerve head (~1000 μm eccentricity). Images were cropped to 150 μm^2^ and RBPMS+ cells and DAPI nuclei counted using the cell counter plugin for Fiji ((57); only round nuclei were counted, thus excluding vascular endothelium); counts were expressed as a density per 100 μm^2^, averaged across the 6 images. For measurement of nuclear diameter, 30 nuclei per cropped image were measured using the line tool, giving an average diameter; this was averaged across the 6 images to produce a final average diameter per retina. These analyses were performed on rat retina (n = 10 eyes from NT, 10 eyes from OHT, 9 eyes from OHT-NAM [200mg/kg/d], 12 eyes from OHT-NAM [400mg/kg/d], 12 eyes from OHT-NAM [800mg/kg/d]), C57BL/6J mouse retina from retinal explants (n = 6 eyes for all groups: 0 days *ex vivo* (DEV; control), 0.5 DEV, 1 DEV, 3 DEV, 5 DEV, 3 DEV + NAM [100mM], 3 DEV + NAM [500mM]), C57BL/6J mouse retina from intravitreal rotenone injection (n = 8 eyes from DMSO control, 8 eyes from rotenone injected, 8 eyes from rotenone + NAM [500mg/kg/d]). Axon integrity in the nerve fiber layer (NFL) of retina following axotomy explant was assessed relative to control (n = 6 eyes for all groups: 0 days *ex vivo* (DEV; control), 3 DEV, 3 DEV + NAM [100mM], 3 DEV + NAM [500mM]). Images of β-tubulin labelled axons adjacent to the optic nerve head were acquired on a Zeiss Axioskop 2 plus epifluorescence microscope, cropped to 250 x 50 μm size and the size and number of varicosities measured. Images were converted to 8 bit, thresholded (200-255) and binarised. Varicosities were analyzed using the ImageJ particle analyzer (FIJI; 1-10 μm size filter to remove single pixel and long, thin axon segments (58)). The number of particles / mm^2^ and the average size of particles (μm^2^) were taken as varicosity density and size. Dendritic degeneration was assessed in axotomy explant retina through DiOlistic labelling of individual RGC dendritic arbors (using a Helios gene gun system, Biorad, as previously described ((51))). Bullets were prepared at a ratio of 2mg DiI, 4mg of DiO, and 80mg Tungsten (1.7 μm diameter) to 30.5 cm of Tezfel tubing (Biorad), by dissolving the dyes in 400 μl of methylene chloride and applying over the tungsten to air dry. The coted tungsten was collected and transferred into the tubing, and distributed along the length of the tubing by vortexing. Individual bullets were cut to 1.2 cm. C57BL/6J mouse retinas were subject to axotomy explant (n = 10 eyes for 0 days *ex vivo* (DEV; control), 12 eyes for all 3 DEV groups: 3 DEV, 3 DEV + NAM [100mM], 3 DEV + NAM [500mM]) and labelled ballistically. For 0 DEV, retinas were dissected in HBSS and immediately labelled, for 3 DEV groups retinas were removed from culture and labelled. For labelling, retinas were flat mounted on to glass slides, all liquid was removed, a culture insert was inverted over the retina to act as a filter to large particle clumps, and the contents of a single bullet discharged at a gene gun pressure of 120 psi. Tissue was transferred to a culture dish containing Neurobasal-A media (supplemented as in the retina axotomy explant model above) and maintained at 37°C, 5% CO_2_ for 30 mins. Retinas were then fixed in 4% PFA for 1 hr, washed in 1M PBS, nuclei labelled with Hoechst (Invitrogen) for 15 mins, washed again and mounted with FluorSave reagent (Merck). After drying, coverslips were sealed. Images of individual RGC dendritic arbors were acquired on a Leica SP8 confocal microscope (Leica Systems; 20X magnification, 0.45 μm/pixel, 1 μm *z*-thickness). Whole dendritic arbors were reconstructed using Imaris (version 9.3.1, Bitplane) where individual RGC dendritic fields were manually selected as areas of interest (used to calculate dendritic field areas) and dendrites automatically traced using the filaments tool. Total dendritic length was calculated and Sholl analysis performed (dendritic intersections per binned distance from the soma center, 10 μm steps) and area under the curve (AUC) of the Sholl curve was calculated. Dendritic health and integrity was evaluated by quantifying dendrite varicosities. Varicosities were identified automatically by the spot tool and a density per cell calculated by normalizing to total dendritic length and dendritic field area.

### Mitochondrial RNA to nuclear RNA ratio

The ratio of RNA from genes encoded in the mitochondrial (mt) and nuclear (nu) genome can differential expression of mitochondrial genes or loss/gain of mitochondrial numbers (59). Six C57BL/6J mouse retina following 3 days ex vivo of the retina axotomy explant model, and six 0 DEV controls were homogenized into 400 μl buffer RLT (Qiagen) with 1% β-mercaptoethanol (Fisher Scientific) using a QIAshredder kit (Qiagen) according to the manufacturer’s instructions. RNA was extracted using RNeasy Mini Kits (Qiagen) according to the manufacturer’s instructions. RNA was extracted into nuclease-free water, and RNA concentration was measured in a 1 μl sample diluted 1:200 in nuclease-free water in a spectrophotometer (BioPhotometer, Eppendorf). cDNA was synthesized using 1 μg of input RNA with an iScript™ cDNA Synthesis Kit and MyIQ thermocycler (both Bio-Rad) and stored at −20 °C overnight. RT-qPCR was performed using 1 μg of input cDNA, 7.5 μl of SsoAdvanced Universal SYBR Green Supermix and 1 μl of the following DNA templates (Prime PCR Assay, Bio-Rad): *mt-Co2* (mitochondrial; *mus musculus), Rps18* (nuclear; *mus musculus*), and *Tbp* (housekeeping; *mus musculus*). A MyIQ thermocycler was used with a 3 min activation and denaturation step at 95 °C, followed by an amplification stage comprising 50 cycles of a 15 sec denaturation at 95 °C and 1 min annealing and plate read at 60 °C. Analysis was performed according to the ΔΔCT method outlined by Quiros *et al*. (59) giving a mtRNA:nuRNA ratio. ΔΔCTs were compared by student’s *t*-test.

### Metabolomics

BN rats were induced with OHT for 3 days or remained as NT controls (as described above). Rats were either pretreated with NAM (800 mg/kg/d) for 7 days (10 days total exposure) or were untreated (n = 8 eyes NT, n = 8 eyes OHT, n = 8 eyes NT-NAM, n = 8 eyes OHT-NAM). Rats were euthanized by pentobarbital injection and cervical dislocation (as described above). Retinas, were immediately dissected on ice cold HBSS (optic nerve head was not included), wiped dry and weighed before snap freezing on dry ice. A second investigator removed the brain and attached optic nerves immediately after death. Optic nerves were separated at the chiasm and weighed before freezing as for retinas. The superior colliculus (SC) and dorsal lateral geniculate nucleus (dLGN) were isolated and separated into left and right hemispheres before weighing and freezing as for the retina. Tissue was stored on dry ice and shipped to the Swedish Metabolomics Centre for sample processing. 200 μl of 90:10 MeOH:H2O was added to each frozen sample on dry ice, acid washed glass-beads (425-600 μm; Sigma Aldrich) were added to constitute 50% v/v of the MeOH:H2O solution and tissues were disrupted by shaking at 30 Hz for 3 mins, followed by an additional shaking step after the addition of 100 μl of H2O. Samples were then centrifuged (14,000 RPM, 10 mins, 4 °C) and 100 μl of the supernatant was transferred to HPLC vials and immediately analyzed. 2 μl was injected into an Agilent 1290 UPLC-system connected to an Agilent 6550 Q-TOF mass spectrometer with an Agilent Jet Stream electrospray ionization (ESI) source. Data was collected in positive and negative ionization mode. The HPLC column used was an HILIC (iHILIC-Fusion(+), 100 x 2.1 mm, 3.5 μM, 100 Å, Hilicon AB). HILIC elution solvents used were A) 50 mM ammonium formate in H2O and B) 90:10 Acetonitrile:[50 mM ammonium formate in H2O]. Chromatographic separation at a flow rate 0.4 mL/min was achieved using the following linear gradient: min 0 = 90% B, min 4 = 85% B, min 5 = 70% B, min 7 = 55% B, min 10 = 20% B, min 10.01 = 90% B, min 15 = 90% B. Seventy three low molecular weight metabolites that could be verified with standards were detected. Metabolites were quantified as area under the curve of the mass spectrometry peak and normalized to an internal standard for negative and positive runs, then normalized to tissue weight. Data were analyzed using MetaboAnalyst (version 4.0, (60, 61)). Groups were compared by two-sample *t*-tests with an adjusted *P* value (false discovery rate, FDR) cutoff of 0.05 deemed significant. Dendograms were created using hierarchical clustering (HC) in R (1-cor, Spearman’s *rho*). Heatmaps were created in R and using Morpheus (https://software.broadinstitute.org/morpheus). Principle component analysis was calculated using Pareto scaling to reduce the effect of magnitude (62), without excluding this dimension entirely. Pathways analysis was performed in MetaboAnalyst using the *Rattus norvegicus* KEGG library. Circos plots were created in R using the *circlize* package (63), using Spearman’s rank correlations with a *P* value adjusted for multiple comparisons.

### Luminometry-based NAD assays

To measure NAD concentration in retina and optic nerves following 14 days of OHT, BN rats were anaesthetized by intraperitoneal injection of pentobarbital (75 mg/kg), and euthanized by cervical dislocation (n = 6 eyes and optic nerves from NT, 6 eyes and optic nerves from OHT). Whole eyes were removed and retina dissected in HBSS as described above. The brain was removed with attached optic nerves. Optic nerves were cut at the chiasm and 4mm from the end proximal to the eye was measured and collected and stored at 4°C in HBSS until further processing. Retinas were transferred to 500 μl of dispase (5000 U, Corning) and put on a heating block (Thermomixer C, Eppendorf) at 37 °C, 350 rpm for 30 mins before dissociation by gentle trituration. Cell concentration was determined by cell counting on a hemocytometer (C-Chip, NanoEntek) and each sample was diluted to a concentration of 2 million cells / ml. Retina and optic nerve samples were then homogenized for 20 secs, 30000 min^-1^ (VDI 12, VWR). NAD concentration was performed using a bioluminescent assay (NAD/NADH Glo-™, Promega). Kit reagents were prepared according to the manufacturer’s instructions. NAD-standards were prepared in HBSS from a 2 mM NAD stock (no. N8285, Sigma-Aldrich). Either 50 μl of sample (100,000 cells / well for retina) or NAD standard were combined with 50 μl of reagent in a 96-well plate (Nunc™ F96 MicroWell™ White Polystyrene plate, ThermoFisher Scientific). Luminescence was recorded at approx. 1 hour from initial mixing (plate shake in machine) to give sufficient time for the reaction to occur and remain stable. Luminescent intensity was converted to NAD concentration using the NAD standard curve. In order to assess how quickly nicotinamide could be utilized by cells to increase NAD concentration NAD assays were performed on treated retina, optic nerve, and brain cortex. C57BL/6J mice (n = 4 mice) were euthanized by cervical dislocation and eyes and optic nerve dissected as above with the exceptions that retinas from left and right eyes were pooled to form a single sample (giving a concentration after processing of 1.4 million cells / ml) and where optic nerves were cut to 2 mm. Whole cortex was removed, separated by hemisphere, transferred to 800 μl of dispase and processed as for the retina above to achieve a concentration of 2 million cells / ml. From each sample (retina, optic nerve and cortex) 4 aliquots of cell homogenate were taken for incubation with NAM (0 mM, 0.1 mM, 10 mM) for either 0, 2, 4, or 6 hours. NAM was added to the correct concentration to the samples so that all incubation times would complete at the same time point. The bioluminescent assay was then conducted as above.

### Generation and Characterization of MitoV mouse

MitoV mice were an independent founder (substrain 1819) in the generation of transgenic mice described by Misgeld *et al*. and as detailed by Burgess and Fuerst (64, 65). YFP is expressed under a rat *Eno2* promoter (neuron specific) and localized to mitochondria via a Cox8a gene-targeting signal fused to the YFP N-terminus. Mice are on a C57BL/6J background. We selected this line based on specificity of inner retinal expression (designated MitoV for visual tissue). Four week old mice were euthanized by cervical dislocation, eyes were enucleated and fixed in PFA for 1 hour, before performing immunofluorescent labelling as described above with anti-GFP and anti-RBPMS antibodies. Brains were fixed in PFA and processed for whole brain cryo-sectioning as described above.

### High resolution fluorescent imaging of mitochondria and morphological analysis

High resolution fluorescent images of mitochondria were captured using Airyscan imaging on a Zeiss LSM800-Airy (63X, 1.3X optical zoom, 66.76 x 66.76 μm images, 35 nm / pixel, *z*-stacks with 15 nm slice thickness). For MitoV mouse tissue, images were acquired of Alexa Fluor 488 conjugated secondary antibodies targeting anti-GFP primary antibody to limit loss of signal from potential YFP bleaching. To allow for practical acquisition times, only the mitochondrial channel was imaged; DAPI nuclear stain and RBPMS labelling in the same image window were captured as snapshots only for reference purposes. Secondary antibody only controls were used for all experiments to set suitable imaging parameters. Retina axotomy explant was performed on MitoV retinas, with 4 retinas maintained for 0 DEV (control), 4 retinas for 0.5 DEV, and 4 retinas for 0.5 DEV with NAM supplemented media (500 mM). Retinas were imaged as flat mounts, with 4 images collected at 1000 μm eccentricity from the optic nerve head superiorly, nasally, inferiorly, and temporally. Images were captured as *z*-stacks from the NFL to the lower boundary of the GCL. The intravitreal rotenone injection model was performed with 9 DMSO (vehicle only) injected retinas, 4 Rotenone (10 mM) injected retinas and 7 Rotenone injected retinas where animals were pre-treated with NAM (500 mg/kg/d) for 1 week prior to injection and maintained for the duration of the experiment. Eyes were enucleated 24 hrs after injection, fixed in PFA and processed for cryo-sectioning as described above. 20 m cryosections were cut and immunofluorescent labelling performed using Alexa Fluor 488 conjugated secondary antibodies targeting anti-GFP primary antibody. Airyscan images were acquired of central retina ~500 μm later to the optic nerve head, encompassing the NFL, GCL and inner plexiform layer (IPL). The IPL was cropped for analysis. In rat tissue, 5 NT eyes (from 3 rats), 6 OHT eyes (from 4 rats) and 4 OHT-NAM (from 4 rats, pre-treatment with 800 mg/kg/d NAM) were collected 14 days after OHT induction and processed for cryo-sectioning as described above. Alexa Fluor 568 conjugated secondary antibodies targeting anti-TOMM20 primary antibodies was imaged. Image capture and analysis followed the same protocol as for rotenone injected eyes. IPL and GCL/NFL crops were analyzed separately. In all three models, mitochondria were reconstructed in 3D using Imaris software (version 9.3.1). Volume reconstructions were performed using the surfaces tool and individual volume and surface area were calculated. Volumes < 125 voxels were filtered and removed from subsequent analysis to reduce noise.

### Live-imaging of mitochondria

RGC cultures were established according as previously described ((66), modified from (67)). Six to seven day old C57BL/6J pups were euthanized and retinas dissected (*n* = 16-18 per panning procedure, 3X repeats). Retinas were dissociated in DPBS (Thermofisher, UK) by forceps within 35 minutes at room temperature under 4X dissecting microscope followed by gentle trituration. RGCs were purified using two-step immune-panning on two Lectin coated negative panning plates (Bandeiraea simplicifolia, BSL-1; Vector Laboratories Ltd.) and a Thy1.2 antibody coated positive plate (Serotec MCA02R, Bio-Rad), over 45 mins at room temperature (25°C). RGCs were seeded at a density of 50,000 cells per well (24 well plate; Nunc, Thermofisher) on Poly-D-lysine and laminin coated glass coverslips (Sigma-Aldrich, R&D Systems, and Academy Science Limited, respectively). RGCs were cultured in serum-free growth medium (37°C, 10% CO_2_) for 10 wells as previously described and treated with NAM dissolved in the media at concentrations of 0, 50, 100, 500, and 1000 uM at day 7 post seeding. At eight days post seeding, mitochondria were stained with MitoTracker Red CMXRos (Molecular Probes, Invitrogen) for 30 mins at 37°C. Using a Zeiss LSM 880 confocal microscope (63X oil immersion lens, 0.132 μm / px) time-lapse imaging was performed in order to record mitochondrial movement. Images were captured every five seconds for 10 minutes. Mitochondria were classified as either stationary or moving over the 10 minute recording based on a total 5 μm distance threshold. Mitochondrial length (long axis), and number of mitochondria (total, motile, stationary) in the entire length of neurites in each group was measured in the first frame (FIJI) and Kymographs of mitochondrial movements were generated using the KymoAnlyzer plugin (68) for FIJI. Mitochondrial velocity was calculated as distance travelled on the kymograph *x* axis divided by length of the *y* axis (duration of movement).

### Extracellular acidification rate and oxygen consumption rate

Retinal ganglion cells (RGCs) were purified from mice according to the methods previously described by Skytt *et al*. and Vohra *et al*. (69, 70). Murine RGCs were cultured in a Seahorse 96-well cell culture microplate at a density corresponding to 450.000 cells / cm^2^. Glycolytic and mitochondrial respiratory function was measured using real-time assessment of, respectively, the extracellular acidification rate (ECAR) and oxygen consumption rate (OCR) with the Seahorse XFe96 Extracellular Flux Analyzer (Seahorse Biosciences-Agilent Technologies, USA) after one week. On the day prior to the assay, cells were pre-incubated 24 hours before the analysis under the different experimental conditions: Control (0 uM nicotinamide (NAM)), 50 uM NAM, 500 uM NAM and 1000 uM NAM. Prior to the assay, culture media was changed to unbuffered DMEM (pH 7.4) and the cells were equilibrated for 15 min at 37°C in a CO_2_-free incubator. Time of calibration inside the seahorse instrument was set for further 15 minutes. Each OCR measurement cycle consisted of 3-min mix and 3-min measurement of the oxygen level, as well as ECR. The analysis of glycolytic and mitochondrial function was initiated by three baseline OCR measurement cycles. The pH of the reagents used to test mitochondrial function was adjusted to 7.4. The OCR and ECR were recorded and calculated by the Seahorse XFe96 software, Wave (Seahorse Biosciences-Agilent Technologies, USA). To normalize the data after the Seahorse analysis, the protein content for each well was measured using a BCA assay with BSA as standard. Measurements were acquired for 3-5 different cell batches per condition.

### Whole-cell patch-clamp electrophysiology

C57BL/6J mice (12 weeks) were anesthetized with 2 % isoflurane and cervically dislocated. Eyes were enucleated and placed into cutting solution consisting of (mM): 125 Choline-Cl, 2.5 KCl, 0.4 CaCl_2_, 6 MgCl_2_, 1.25 NaH2PO4, 26 NaHCO_3_, 20 D-glucose, on ice and saturated with 95% O_2_ plus 5% CO_2_. The cornea was pierced with a 30 G needle and the retina dissected out. The retina was then incubated for 2 hours at room temperature in Ames medium (Sigma Aldrich; AUS) and 1.9g sodium bicarbonate (Sigma Aldrich; AUS) with either 1 mM, 10 mM or 100 mM, or 250 mM NAM (Sigma Aldrich; AUS) or control solution saturated with 95% O_2_ and 5% CO_2_. Retina were transferred to a submerged recording chamber on an upright microscope (Slicescope Pro 1000; Scientifica, UK). For all recordings, retina were perfused (2 ml/min) with Ames medium saturated with 95% O_2_ plus 5% CO_2_ at 24 °C, containing either 1 mM, 10 mM,100 mM or 250 mM NAM or control (0 mM) solution as per the previous incubation step. The center of the retina was located using infrared-oblique illumination microscopy with a 4x air objective (Olympus, Japan) and a CCD camera (IEEE 1394; Foculus, Germany) and retinal ganglion cells visualized using a 40× water-immersion objective (Olympus, Japan). Retinal ganglion cells with large cell bodies were chosen and their identity confirmed by the presence of AP firing. Patch pipettes (6-9 MΩ; GC150F-7.5; Harvard Instruments; USA) were pulled using a Flaming/brown micropipette puller (Model P-1000; Sutter instruments, USA) and filled with solution containing (mM): 115 K-gluconate, 5 KCl, 5 EGTA, 10 HEPES, 4 Mg-ATP, 0.3 Na-GTP, and 7 phosphocreatine (pH 7.3). Once wholecell configuration was obtained, a holding current was injected to maintain a holding potential of approximately −60 mV. Cells were excluded if a holding current greater than −100 pA was required to achieve a membrane potential of −60 mV. Patch clamp recordings were performed using a PatchStar micromanipulator (Scientifica, UK) and Axon Multiclamp 700B patch-clamp amplifier (MDS, USA). Current clamp data were acquired using pClamp software (v10; MDS, USA) with a sampling rate of 100 kHz and low pass Bessel filtered at 10 kHz (Digidata 1440a; Axon). Action potential firing was recorded using an episodic protocol with the following current steps (0 to 200 pA in 10pA steps, 400 ms duration, 5s inter sweep interval).

### Analysis of action potential firing

Data were analyzed using AxoGraph X software (Berkeley, CA, USA). Action potentials were detected using an amplitude threshold of 50 mV relative to pre-event baseline. Input-output curves were calculated by determining the number of action potentials generated for each injected current step. All action potential kinetics were analyzed for the first action potential at rheobase current, defined as the minimum current injection required to generate an action potential. Action potential threshold was defined as 30 mV/s. Action potential amplitude was calculated relative to pre-event baseline. Action potential half-width was calculated at 50% of maximal action potential amplitude. The action potential rise-time was calculated as the time between 20% and 80% of the maximal action potential amplitude.

### Statistical analysis

All statistical analysis was performed in R. Data were tested for normality with a *Shapiro Wilk* test. Normally distributed data were analyzed by *Student’s t*-test or *ANOVA* (with *Tukey’s HSD*). Non-normally distributed data were transformed using squared transforms; data that remained non-normally distributed was analyzed by a Kruskal-Wallis test followed Dunn’s tests with Benjamini and Hochberg correction. Unless otherwise stated, * = *P* < 0.05, ** = *P* < 0.01, *** *P* < 0.001, NS = non-significant (*P* > 0.05). For box plots, the center hinge represents the mean with upper and lower hinges representing the first and third quartiles; whiskers represent 1.5 times the interquartile range.

## Supplementary Materials

The following supplementary materials are attached:

Supplementary Tables 1-3

Supplementary Figures 1-4

## Acknowledgments

The Authors would like to thank Monica Aronsson and Diana Rydholm for their assistance with animal husbandry and maintenance, St. Eriks Eye Hospital for financial support for research space, clinical histopathology, and animal facilities, the Simon John Lab at The Jackson Laboratory for assistance with breeding and shipping animals, the Knut and Alice Wallenberg Foundation and Karolinska Institutet for supporting the CLICK imaging facility, and the Swedish Metabolomics Centre for assistance developing the small molecular weight metabolomics.

## Author contributions

JRT – designed and performed experiments, analyzed data, wrote the manuscript; AO – performed experiments, analyzed data; SS – performed experiments, analyzed data; SAE – performed experiments, analyzed data; GC – performed experiments, analyzed data; RV – performed experiments, analyzed data; MJ – performed experiments, analyzed data; EL – performed experiments; APV – analyzed data; AD-V – analyzed data; EK – performed experiments; SR – analyzed data; GJ – provided resources and expertise, designed experiments; PGF – provided resources and expertise, designed experiments; RWB – provided resources and expertise, designed experiments; RB – provided resources and expertise, designed experiments; MK – provided resources, designed experiments; JEM – provided resources, designed experiments; JGC – provided resources, designed experiments; MV – provided resources, designed experiments; PAW – conceived, designed, performed experiments, analyzed data, wrote the manuscript. All authors read and approved the final manuscript.

---

**Supplementary Figure 1. Retinal axotomy explant model**. (**A**) Eye enucleation severs the optic nerve, resulting in significant RGC death (loss of RBPMS+ cell somas; an RGC specific marker in the retina, red), as the retina is maintained in culture (*n* = 6 retinas per condition). (**B**) A significant reduction in RGC density, nuclear density, and nuclear diameter (DAPI, blue in A) occurs as early at 0.5 days ex vivo (DEV) and is highly significant at 1, 3, and 5 DEV. (**C**) β-tubulin labelling (green) and imaging of the retinal nerve fiber layer demonstrated (**D**) a significant increase in axon varicosity density by 3 DEV (but not in varicosity size; *n* = 6 retinas per condition). Scale bar = 20 μM in B and 50 μm in B. ** *P* < 0.01, *** *P* < 0.001.

**Supplementary Figure 2. Low molecular weight Metabolomics.** (**A**) Seventy nine low molecular weight metabolites could be reliably detected in retina (*n* = 8), optic nerve (*n* = 8) and superior colliculus (SC; *n* = 8 hemispheres) of Brown Norway rats. Unsupervised hierarchical clustering clearly separated these tissue based on their metabolic profile (accompanying heat map scaled where red = highest value, blue = lowest value by row). The majority of metabolites were most abundant in the optic nerve and lowest in the retina (with the exception of ATP; greyscale from highest (black) to lowest (light grey)). (**B**) A number of metabolites were highly abundant and so data were subject to Pareto scaling in order to reduce this as a driver for inter tissue differences (metabolites match labels on A). (**C**) These tissue differences were explored by principle component analysis. Metabolite abundance per sample (relative area) for the 5 explanatory variables with the greatest loadings for components 1 and 2 (PC1 and PC2) are plotted demonstrating inter tissue differences in these metabolites.

**Supplementary Figure 3. NAD metabolism is altered under NAM treatment and OHT across the retina, optic nerve and superior colliculus.** Metabolite abundance per sample (relative area) for NAD, NADH, and Nicotinamide (NAM) are plotted demonstrating differences in these metabolites by condition (NT, NT-NAM, OHT, OHT-NAM; *n* = 8 samples for each tissue and condition). The NAD:NADH ratio is also plotted. (**A**) In the retina, NAD increases under OHT, and increases under NAM supplementation in both NT and OHT conditions. NADH, NAM and NAD:NADH show little change across conditions in the retina. (**B**) In the optic nerve, NAD, NADH, NAM and the NAD:NADH ratio are significantly increased under NAM supplementation, and remain higher in OHT-NAM. NAM is highly increased in OHT-NAM treated optic nerves, suggesting a potential reduction in conversion to NAD. (**C**) In the superior colliculus NAD is decreased in OHT and this is rescued under NAM supplementation (OHT-NAM). NAD and NADH are significantly increased under NAM supplementation and NAM is unchanged. The NAD:NADH ratio is decreased in OHT and this is rescued by NAM supplementation (OHT-NAM). * *P* < 0.05, ** *P* < 0.01, *** *P* < 0.001, NS *P* > 0.05.

**Supplementary Figure 4. Characterization of MitoV mouse**. In substrain 1819 generated as described by Misgeld *at al*. (2007) YFP is expressed under a rat *Eno2* promoter (neuron specific) and localized to mitochondria via a *Cox8a* gene-targeting signal fused to the YFP N-terminus. (**A**) In the retina YFP expression is visible within RGC soma, axons and dendrites. (**B**) Labelling with an anti-GFP antibody to boost detection of YFP confirmed that 100% of YFP positive cells were all also RBPMS positive (data not shown). (**C**) Retinal sections demonstrate that YFP is also present within a subset of bipolar cells and some photoreceptor outer segments (left *panel*) as demonstrated by areas of non-localization of TOMM20 (all mitochondria) and YFP (*right panel*). (**D**) YFP expression was also present across multiple brain regions and the spinal cord.

**Supplementary Table 4.**

| Antibody | Target | Host | Dilution | Details |
| --- | --- | --- | --- | --- |
| βIII-tubulin | Microtubules specific to RGCs in the retina | Chicken | 1:250 | Novussbio cat # NB100-1612 |
| RBPMS | RNA-binding protein, RGC specific in the retina | Rabbit | 1:500 | Novusbio cat # NBP2-20112 |
| anti-GFP | XFPs | Chicken | 1:500 | Abcam cat # ab13970 |
| TOMM20 | Translocase of outer mitochondrial membrane 20, all mitochondria in the retina, disease resistant expression | Rabbit | 1:500 | Abcam cat # ab78547 |
| Goat-anti Rabbit AF 568 | Rabbit primary antibody | Goat | 1:500 | Abcam cat # A11011 |
| Goat-anti Chick AF 488 | Chick primary antibody | Goat | 1:500 | Abcam cat # A11039 |

