## Supplementary figures and images for "Nicotinamide provides neuroprotection in glaucoma by protecting against mitochondrial and metabolic dysfunction"

### Supplementary Figure 1

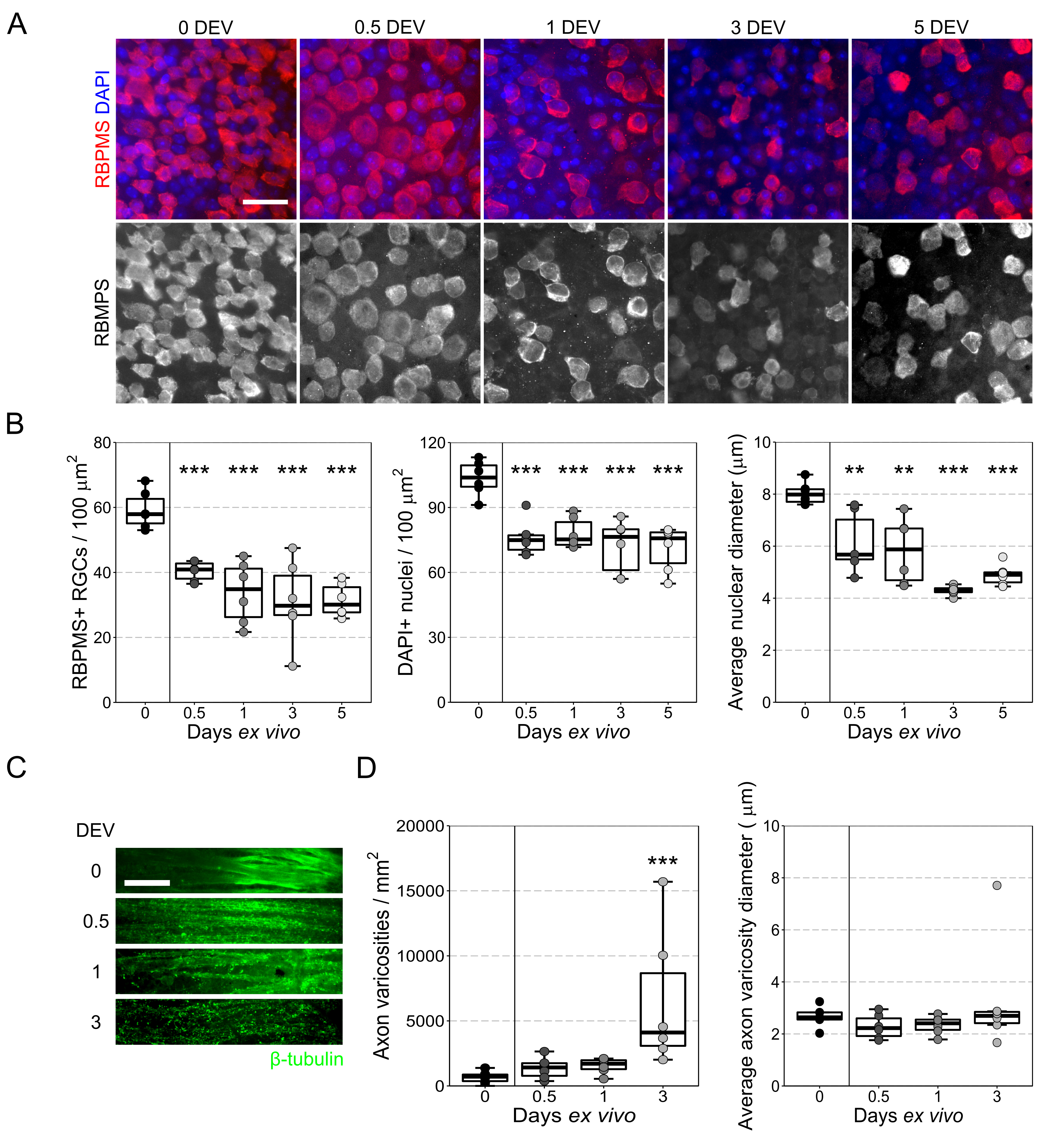

### Supplementary Figure 2

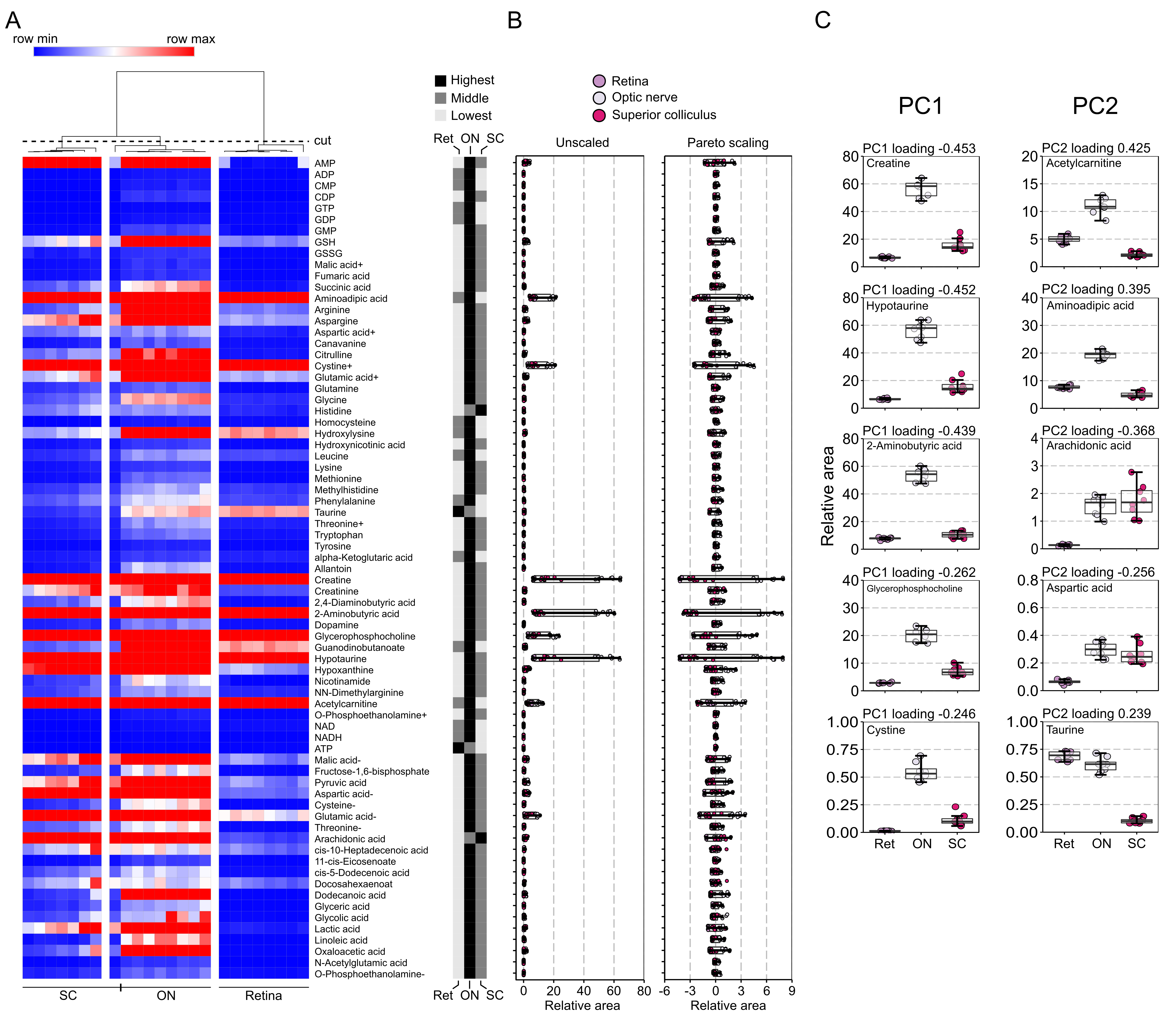

### Supplementary Figure 3

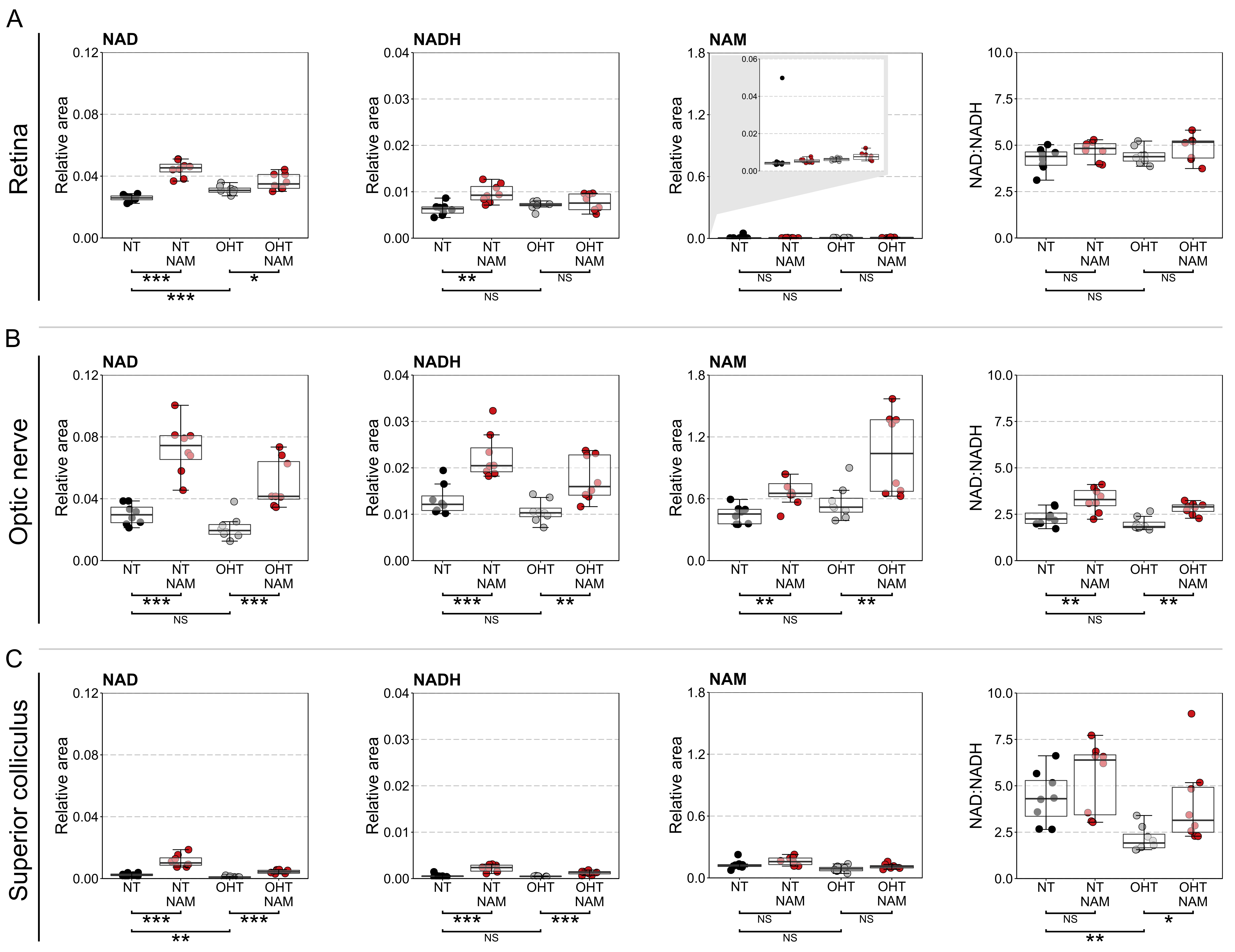

### Supplementary Figure 4

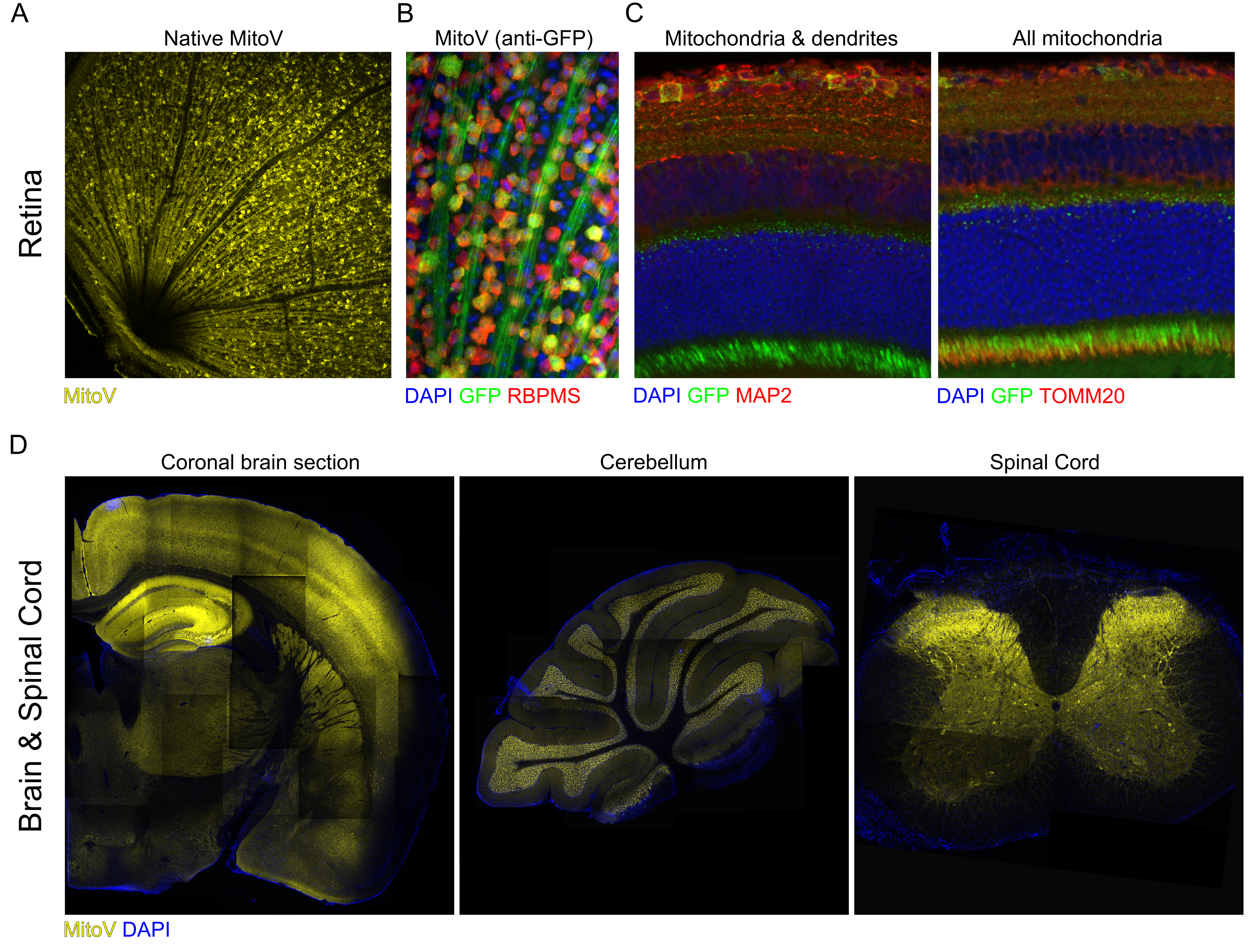
